# Acquisition of ionic copper by a bacterial outer membrane protein

**DOI:** 10.1101/2020.06.04.134395

**Authors:** Satya Prathyusha Bhamidimarri, Tessa R. Young, Muralidharan Shanmugam, Sandra Soderholm, Arnaud Baslé, Dirk Bumann, Bert van den Berg

## Abstract

Copper, while toxic in excess, is an essential micronutrient in all kingdoms of life due to its essential role in the structure and function of many proteins. Proteins mediating ionic copper import have been characterised in detail for eukaryotes, but much less so for prokaryotes. In particular, it is still unclear whether and how Gram-negative bacteria acquire ionic copper. Here we show that *Pseudomonas aeruginosa* OprC is an outer membrane, TonB-dependent transporter that is conserved in many Proteobacteria and which mediates acquisition of both reduced and oxidised ionic copper via an unprecedented CxxxM-HxM metal binding site. Crystal structures of wild type and mutant OprC variants with silver and copper suggest that acquisition of Cu(I) occurs via a surface-exposed “methionine track” leading towards the principal metal binding site. Together with whole-cell copper quantitation and quantitative proteomics in a murine lung infection model, our data identify OprC as an abundant component of bacterial copper biology that may enable copper acquisition under a wide range of conditions.

**Significance:** Copper is an essential metal in biology due to its role in the structure and function of many proteins. Despite this, it is not very clear how bacteria acquire copper, especially for Gram-negative organisms. In this study we show that the outer membrane protein OprC has an unusual metal binding site that allows OprC to bind both reduced and oxidised ionic copper near-irreversibly. Given the versatility of OprC, its presence in many Proteobacteria and its abundance during lung infection in mice, our study shows that OprC is an important component of prokaryote copper biology that warrants further study to uncover its regulation and to assess its role in bacterial virulence.

## Introduction

Metals fulfil cellular functions that cannot be met by organic molecules and are indispensable for the biochemistry of life in all organisms. Copper is the third-most abundant transition metal in biological systems after iron and zinc. It has key roles as structural component of proteins or catalytic cofactor for enzymes(1), most notably associated with the biology of oxygen and in electron transfer. On the other hand, an excess of copper can be deleterious due to its ability to catalyse production of hydroxyl radicals(2, 3). Excessive copper may also disrupt protein structure by interaction with the polypeptide backbone, or via replacement of native metal cofactors from proteins, thus abolishing enzymatic activities via mismetallation (1, 4, 5). Thus, cellular copper levels and availability must be tightly controlled. Bacterial copper homeostasis systems are well characterised(6). Specific protein machineries are involved in fine-tuning the balance of intracellular copper trafficking, storage and efflux according to cellular requirement, in such a way that copper is always bound. This control is executed by periplasmic and cytosolic metalloregulators, which activate transcription of periplasmic multi - copper oxidases, metallochaperones, copper-sequestering proteins(7, 8) and transporters(9–11). To date, relatively few families of integral membrane proteins have been validated as copper transporters, and these have different structures and transport mechanisms(12). The P1B-type ATPases such as CopA are responsible for Cu(I) efflux from the cytosol via several metal binding domains, using energy released from ATP hydrolysis (13–15). A second class of copper export proteins are RND-type tripartite pumps such as CusABC, which efflux Cu(I) by utilising the proton-motive force(16–18). Relatively few copper influx proteins have been identified. The bacterial inner membrane copper importer CcoA is a major facilitator superfamily (MFS)-type transporter involved in fine-tuning the trafficking of copper into the cytosol and required for cytochrome c oxidase maturation(19, 20). The Ctr family of copper transporters is responsible for Cu(I) translocation into the cell without requiring external sources of energy(21). However, Ctr homologs are found only in eukaryotes, and the molecular mechanisms by which copper ions enter Gram-negative bacteria is still a matter of debate. The exception is copper import via metallophores like methanobactin, a small Cu - chelating molecule that is secreted by methanotropic bacteria and most likely taken up via TonB-dependent transporters, analogous to iron-siderophores(22).

*Pseudomonas aeruginosa* is a versatile and ubiquitous Gram-negative bacterium and a notorious opportunistic pathogen in humans, playing a major role in the development of chronic lung infection in cystic fibrosis patients(23, 24). *P. aeruginosa* has a number of TonB-dependent transporters (TBDTs) in the outer membrane (OM) dedicated to the acquisition of different iron-siderophore complexes such as pyochelin and pyoverdin(25). In addition, *P. aeruginosa* contains another TBDT, termed OprC (*PA3790*), whose function has remained enigmatic. Nakae *et al*. suggested that OprC binds Cu(II) with micromolar affinities(26). Transcription of OprC was found to be repressed in the presence of Cu(II) in the external medium under aerobic conditions(26–29), suggesting a role for OprC in copper acquisition. Very recently, the blue copper protein azurin was reported to be secreted by a *P. aeruginosa* Type VI secretion system and to interact with OprC, suggesting a role of the latter in Cu(II) uptake(29).

To clarify the role of OprC in copper biology, we have determined X-ray crystal structures of wild type and mutant OprC proteins in the absence and presence of copper and silver, and characterised metal binding via ICP-MS and EPR. In addition, we have confirmed metal uptake by OprC using whole cell metal quantitation. OprC indeed has the typical structure of a TBDT, and differences between the Cu-loaded and Cu-free protein demonstrate changes in tertiary structure that likely lead to TonB interaction and copper import.

## Results

### OprC is a TonB-dependent transporter that binds ionic copper

The structure of OprC, crystallised with an N-terminal His7 purification tag under aerobic conditions in the presence of 2 mM CuCl2, was solved using single wavelength anomalous dispersion (Cu-SAD), using data to 2.0 Å resolution (Methods; Table S1; Fig. S1). As indicated by the successful structure solution, OprC contains a single bound copper and shows the typical fold of a TBDT, with a large 22-stranded β-barrel occluded by an N-terminal ∼15 kDa plug domain (Fig. 1). The copper binding site comprises residues Cys143 and Met147 in the plug domain and His323 and Met325 in the barrel wall. The CxxxM-HxM configuration, which coordinates the copper in a tetrahedral manner (Fig. 2A, B), is highly unusual and has, to our knowledge, not been observed before in copper homeostasis proteins. A similar site is present for one of the copper ions of the valence-delocalised CuA dimer in cytochrome c oxidase, where the copper ion is coordinated by 2Cys+1Met+1His(30, 31). Other similar sites are class I Type I copper proteins like pseudoazurin and plastocyanin, where copper is coordinated by 2His+1Cys+1Met(32). Interestingly, and unlike class I Type I copper proteins, concentrated solutions and OprC crystals obtained in the presence of Cu (II) are colourless. Another notable feature of the OprC structure becomes apparent when analysing the positions of the methionine residues. As shown in Fig. 2D, out of the 15 visible methionines in OprC, 10 are organised in such a way that they form a distinct “track” leading from the extracellular surface towards the copper binding site. An additional two methionines (Met448, Met558) are not visible due to loop disorder, but given their positions they will be a part of the methionine track. Considering that Cu(II) prefers nitrogen and oxygen as ligands while Cu(I) prefers sulphur, we propose that the methionine track might bind Cu(I) with low affinity and may guide the metal towards the principal binding site, which is at the bottom of the track (Fig. 2D). Importantly, the anomalous difference maps of OprC crystallised with Cu(II) do not show any evidence for weaker, secondary copper sites (Fig. S1), demonstrating that there are no other copper binding sites and that the methionine track indeed does not bind Cu(II).

**Figure 1.**
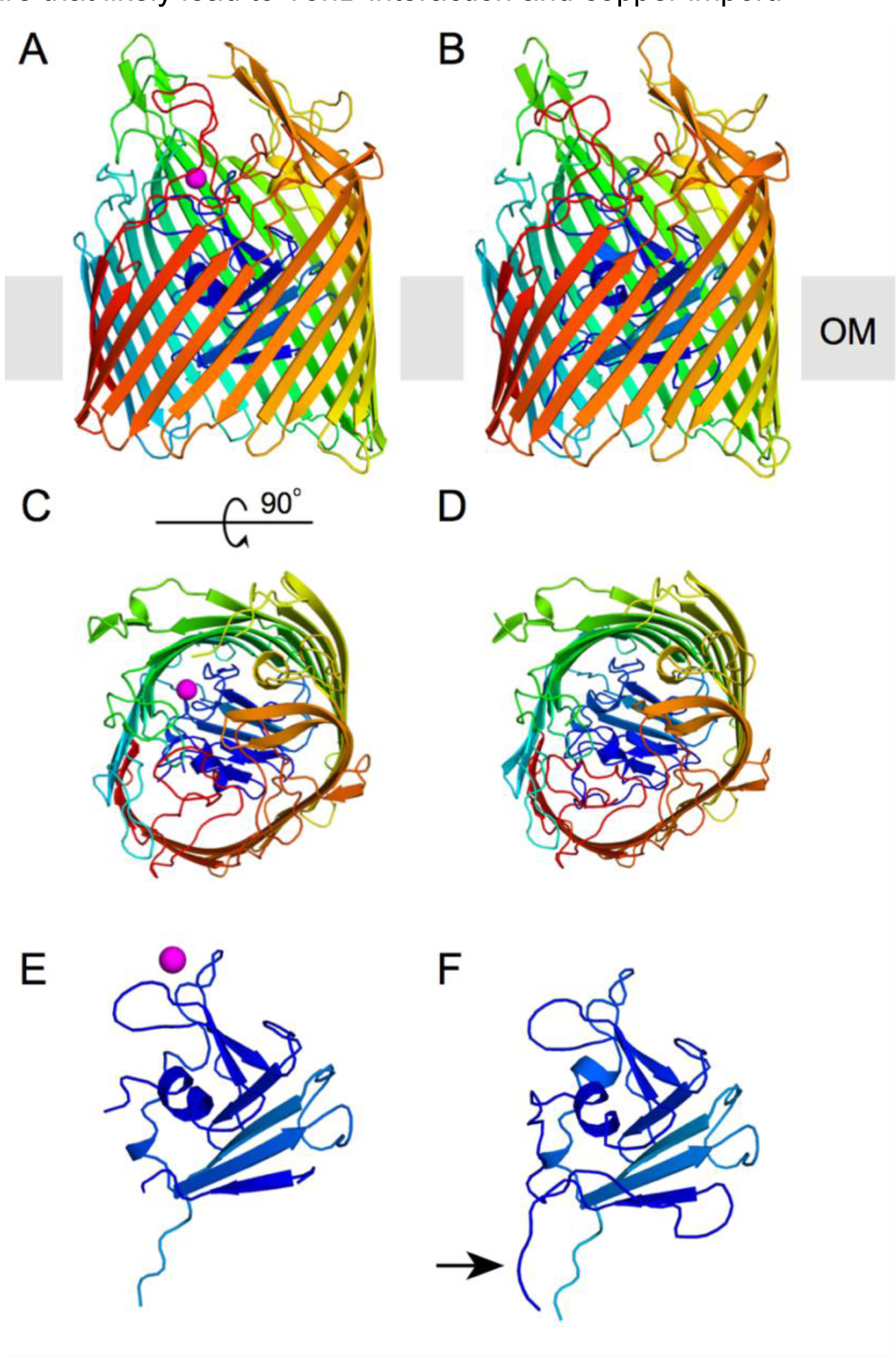
OprC is a TonB-dependent transporter. Cartoon representation of (A,C) Cu-loaded OprC and (B,D) Cu-free OprC (OprC_AA_). The N-terminal plug domain is shown separately for both forms (E,F). Structures are shown in rainbow from N-terminus (blue) to C-terminus (red); copper is represented as a magenta sphere. The arrow in (F) highlights the visibility of the Ton box in apo-OprC.

**Figure 2.**
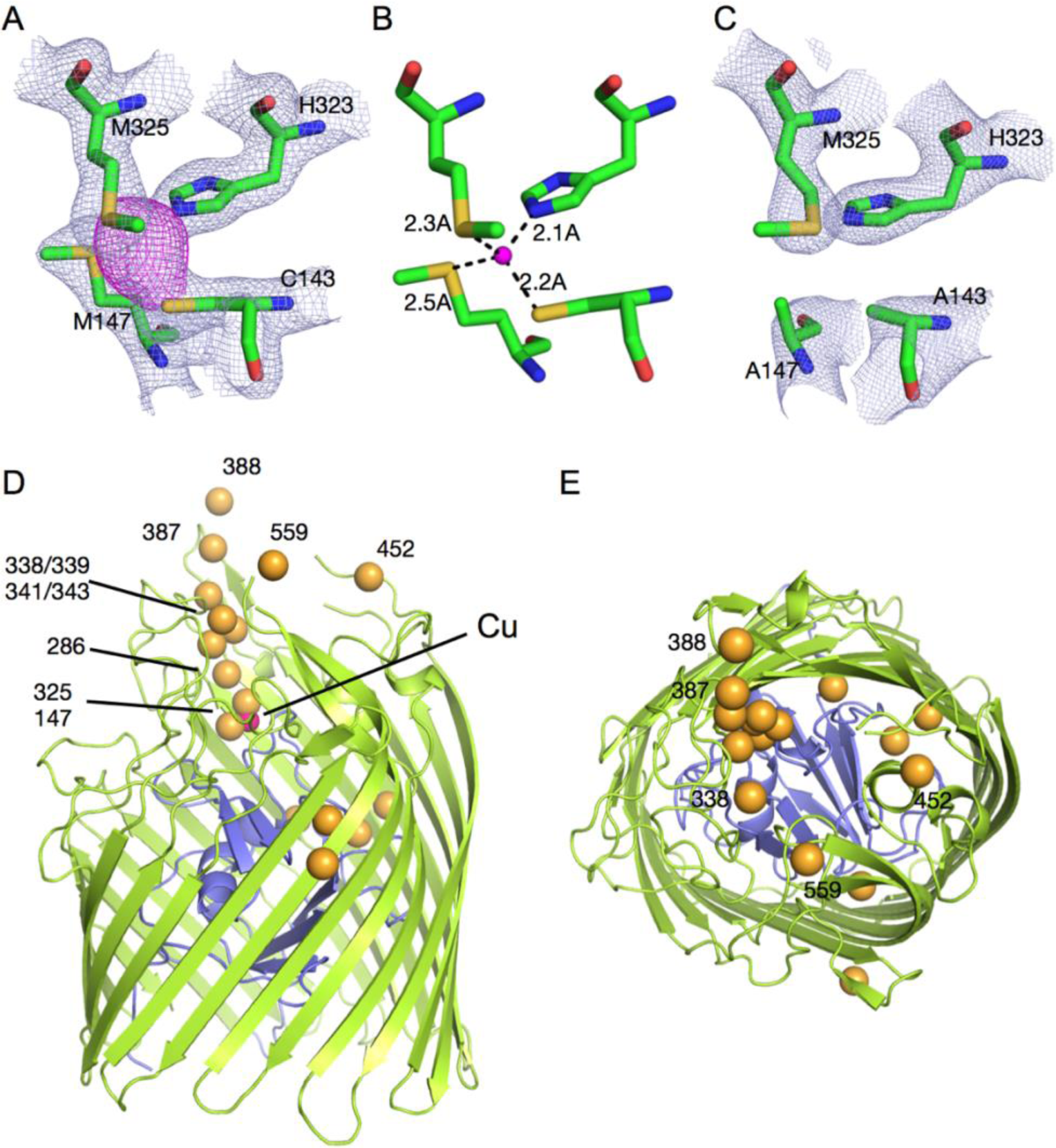
OprC has an unusual CxxxM-HxM binding site and a methionine binding track. (A), Stick models of copper-coordinating residues Cys143, Met147, Met325 and His323. Electron density in gray mesh (2Fo-Fc map contoured at 2.0σ, carve = 2.0) is shown for the binding site residues C/M-H/M and the copper atom (anomalous difference map shown in magenta, contoured at 3.0σ, carve = 2.25). (B) Distances between coordinating residues and metal show that copper is coordinated via 1 thiolate (from Cys), two thioethers (from Met), and one imidazole nitrogen from His. (C) Mutation of binding site residues Cys143 and Met147 to alanines abolishes copper binding (2Fo-Fc map contoured at 2.0σ, carve = 2.0). (D,E) OM plane (D) and extracellular views (E) showing the thioether atoms of all methionine residues present in OprC as yellow spheres. The copper atom, only visible in (D), is shown as a magenta sphere.

### Copper binding by OprC is highly specific and near-irreversible

Following structure determination of copper-bound OprC, several attempts were made to produce a structure of copper-free OprC. First, the protein was purified and crystallised without added copper; however, this gave a structure that was identical to the one already obtained and contained bound copper that presumably originated from the LB medium. As expression in rich media always yielded OprC with 0.5-0.8 equivalents copper as judged by ICP-MS, various attempts to lower the copper content were made. Removal of bound copper from purified protein with combinations of denaturants (up to 4.0 M urea) and EDTA were not successful. Expression in minimal medium reduced, in the best cases, the metal content of the wild type to ∼45 % equivalency (Figs. 3A, B). Subsequent aerobic incubation of OprC for 30 min in the presence of either 3 or 10 equivalents Cu(II) followed by size exclusion chromatography (SEC) in buffer containing 0.5 mM EDTA demonstrate co-elution of 1 equivalent copper (Fig. 3B). Thus, the His7 tag does not bind Cu(II) with high affinity. Co-incubation with 0.5 mM EDTA (∼50-fold excess) does not result in copper loading, suggesting that EDTA effectively withholds Cu (II) from OprC (Fig. S2). As-purified OprC does not contain zinc, the most common contaminant in metal-binding proteins, nor does it contain appreciable amounts of any other metals that could have been introduced during purification such as Ni and Fe, indicating that OprC is highly specific (Fig. 3A, Fig. S2). Indeed, incubation of purified OprC in the presence of 3 or 10 equivalents Zn does not result in zinc co-elution (Fig. S2). To obtain copper-free OprC after purification from rich media, we constructed a variant (OprC_AA_) in which the binding site residues Cys143 and Met147 were both mutated to alanines (Fig. 3A; AA). Even after equilibration of OprC_AA_ for 30 min with 3 or 10 equivalents Cu (II) no co-elution with metal is observed (Fig. 3B), indicating that high-affinity copper binding is completely abolished and confirming that the His7 tag does not bind Cu(II) with an affinity high enough to survive SEC, possibly due to the presence of 0.5 mM EDTA in the SEC buffer.

**Figure 3.**
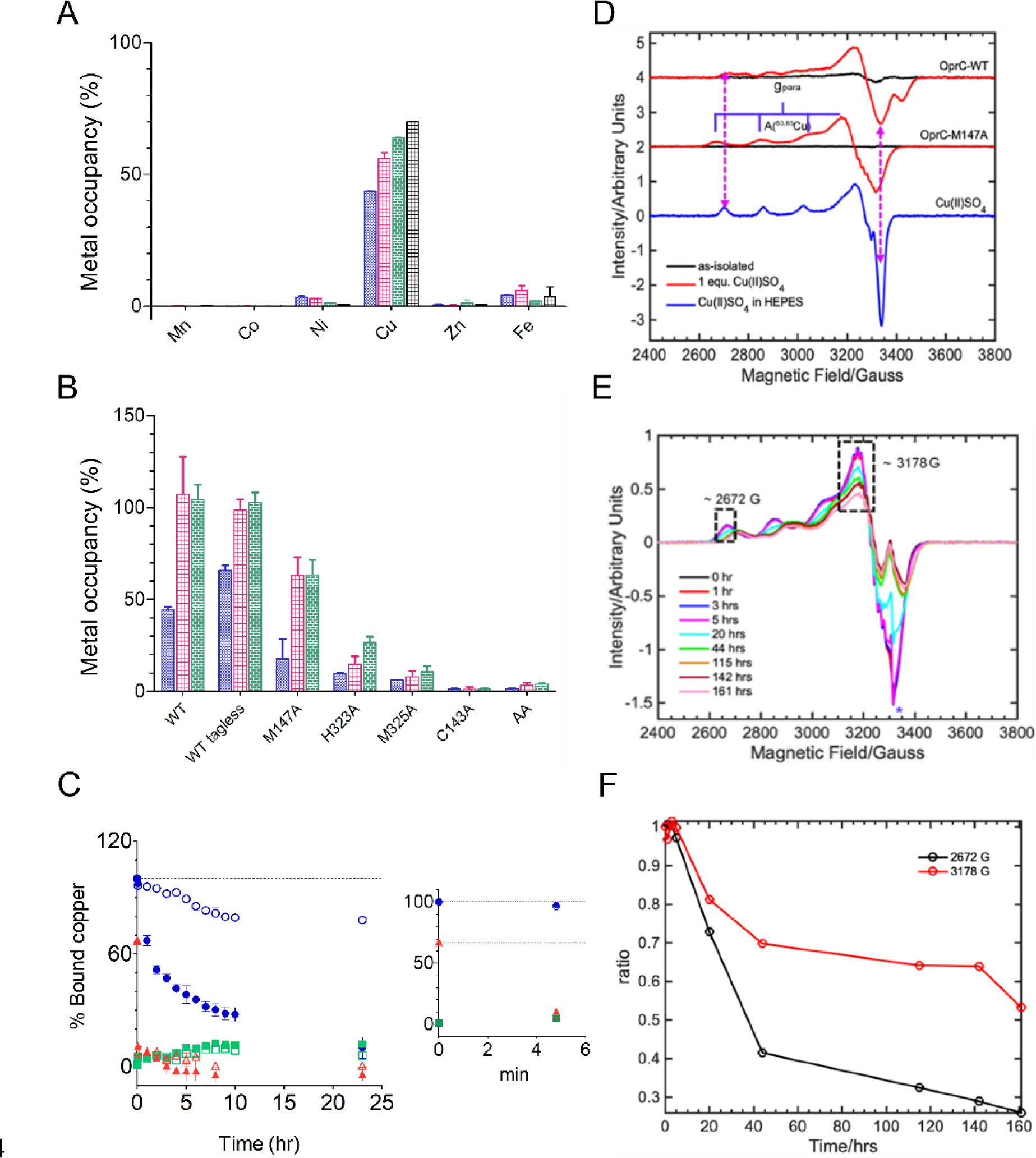
OprC binds 1 equivalent copper near-irreversibly. (A) Metal occupancy of as-purified wild type OprC by ICP-MS shows specific binding only to copper. Each colour indicates an individual batch of purified protein. Averages ± s.d are shown (3 technical replicates) (B) Copper content of wild type OprC and binding site mutant proteins before (blue) and after aerobic incubation with either 3 (pink) or 10 (green) equivalents Cu(II) for 30 min followed by analytical SEC. All proteins except where stated contain a N-terminal His7 tag. Averages ± s.d. of three or four incubations from one or two different protein batches are shown (n = 3 or 4) (C) Copper is kinetically trapped in OprC. Time course of copper extraction experiments showing % bound copper for OprC WT (blue), OprC_AA_ (green) and OprC M147A (red), at room temperature (open symbols) and 60 °C (filled symbols). The inset shows % bound copper in the first few minutes after starting the experiment. OprC_AA_ served as a control. Dotted lines indicate initital occupancies of OprC WT and M147A. (D) Comparison of the cw-EPR spectra of OprC_WT_ and OprCM147A mutant before (black traces) and after (red traces) addition of Cu(II) solution to 1 equivalent. The blue trace shows the EPR spectrum of the Cu(II)SO4 in Hepes buffer. All EPR spectra have been background subtracted. The double-headed magenta dotted arrows show the difference in the observed **g** and **A** tensors of OprC variants. The blue goal-posts indicate the ^63,65^Cu-hyperfine splitting along the parallel region. Before Cu(II) addition, the copper equivalencies were 0.6 for OprC_WT_ and 0.1 for OprCM147A. (E) EPR time course for OprCM147A after addition of 1 equivalent Cu (II). (F) Relative intensities of EPR signals at ∼ 2672 G and 3178 G (black dotted rectangular boxes in the top panel) plotted as a function of time. Values shown are averages from three independent time courses.

The fact that it is not possible to obtain copper-free wild type protein, even after taking extensive precautions, suggests that copper binds to OprC with very high affinity. To explore this further, we performed copper extraction assays with a large excess of bathocuproinedisulfonic acid (BCS) under reducing conditions (Methods). For copper -loaded OprC_WT_, only 20% copper was removed after 24 hrs at room temperature, and the temperature had to be increased to 60 °C to obtain near-quantitative extraction of copper (∼90% after 24 hours) (Fig. 3C). For reasons that are unclear, the orange-coloured [Cu(BCS)2]^-3^ complex was hard to separate from OprC, and BCS-treated OprC did not bind copper anymore, suggesting an irreversible change in the protein due to the harsh incubation conditions. Nevertheless, these results demonstrate that copper is kinetically trapped inside OprC and is, for all intents and purposes, irreversibly bound. This is fully compatible with the consensus transport mechanism of TBDTs, in which the interaction with TonB, occurring after substrate binding, is required to disrupt the binding site, leading to release of the substrate(33).

### Conformational changes upon copper binding

The OprC_AA_ structure was solved by molecular replacement using Cu-bound OprC as the search model (Figs.1 B,D,F; Fig. 2C). The binding site residues of both structures occupy very similar positions, indicating that the introduced mutations abolish copper binding with out generating gross changes in the binding site. Superposition of the structures (Figs. 4 A,B) shows that for the remainder of the transporter, structural changes upon copper binding are confined to the vicinity of the copper binding site, with parts far removed virtually unchanged (overall Cα r.m.s.d ∼ 1.0 Å). The largest change is observed for loop L11, which undergoes an inward-directed motion of ∼ 8.0 Å upon copper binding (Figs. 4 D,E and Fig. S3). A similar inward-directed but smaller change occurs for loop L8. Some loop tips (*e.g*. L4, L5, L6) in OprC_AA_ lack electron density for a limited number of residues, suggesting increased mobility. Overall, the conformational changes of the loops upon copper binding likely decrease the accessibility of the copper binding site. However, the main reason why the bound copper is inaccessible to solvent is that the binding site residues Met147 and Met325, together with Asn145, effectively form a lid on the copper ion in the wild type transporter. In the double mutant, copper becomes solvent accessible due to the absence of the Met147 side chain (Fig. 4D, E and Fig. S3).

**Figure 4.**
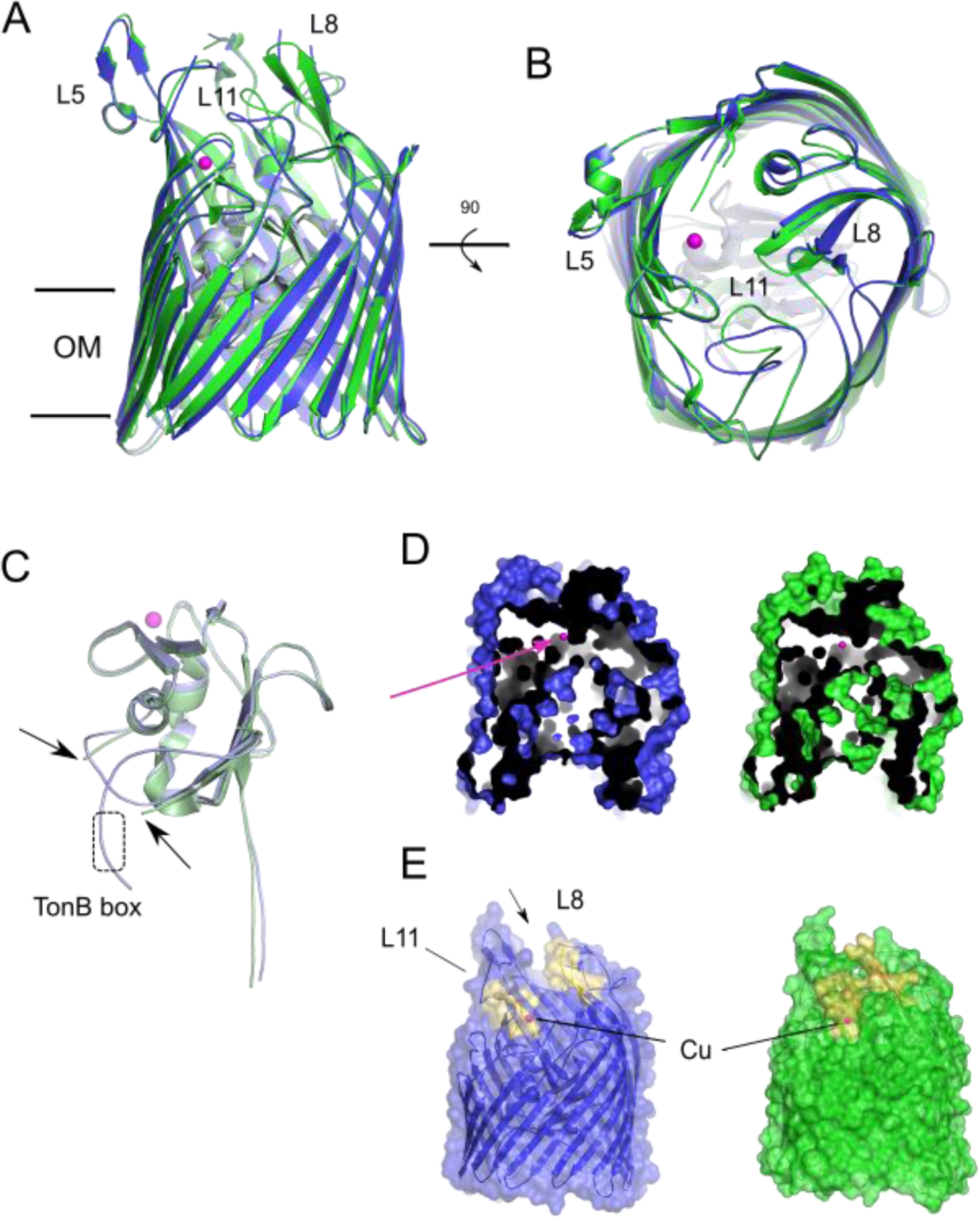
OprC structural changes upon copper binding. (A,B) Cartoon superposition from the OM plane (A) and extracellular environment (B) of OprC_WT_ (coloured green) and OprC_AA_ (blue), indicating locations of loops L5, L8 and L11; copper is represented as a magenta sphere. The plug domains of OprC_WT_ and OprC_AA_ are coloured light green and light blue, respectively. (C) Superposition of N-terminal plug domains indicating the location of the TonB box, which is invisible in Cu-bound OprC_WT_. Arrows indicate the missing density for Glu88-Pro94 in OprC_WT_. (D) Surface slab representations from the OM plane, showing the presence of a solvent pocket in OprC_AA_ that is generated by the absence of Met147 (arrow). For orientation purposes, the OprC_WT_-bound copper is shown in both structures. (E) Side surface views showing the conformational changes of L8 and L11 (coloured yellow) as a result of copper binding. As in (E), the bound copper of OprC_WT_ is shown in both proteins.

The consensus mechanism for TonB-dependent transport postulates that ligand binding on the extracellular side generates conformational changes that are propagated to the periplasmic side of the plug and increase the periplasmic accessibility of the Ton box for subsequent interaction with TonB(34). In OprC_AA_, N-terminal density is visible up to Leu66 (*i.e*., the first 10 residues of the mature protein are disordered) including the Ton box (^68^PSVVTGV^75^), which is tucked away against the plug domain and the barrel wall. In Cu - OprC, the density between Glu88 and Pro94 is hard to interpret and, more importantly, no density is observed before Pro79, including the Ton box (Figs. 1 E,F and 4 C). Thus, while we cannot say conclusively that the Ton box is accessible to TonB in Cu-OprC, the structures do show that changes occur in the Ton box upon substrate binding. Thus, the structures of OprC in the absence and presence of ligand are consistent with the consensus TBDT mechanism. The observed position of the Ton box in OprC_AA_, likely hard to reach from the periplasmic space, would prevent non-productive interactions of TonB with transporters that do not have substrate bound(34).

### The OprC methionine track and the principal binding site bind Cu(I)

We next asked whether OprC also binds Cu (I). Since it is challenging to maintain copper in its +1 state during crystallisation, we used silver (Ag(I)) as a proxy for Cu(I) and determined the co-crystal structure of WT OprC in the presence of 2 mM AgNO3 (Methods). This is possible because the as-purified protein used for crystallisation only had partial copper occupancy (∼60%). Data was collected at 8000 eV, at which energy the anomalous signal of copper is very small (0.6 e^-^, compared to 4.2 e^-^ for Ag). Strikingly, and in sharp contrast to Cu (II) (Fig. S1), the anomalous map of OprC WT crystallised in the presence of silver shows not one but three anomalous peaks. The first, strong peak (Ag1; 23σ) is located at the same site as in OprC crystallised with Cu (II), and is coordinated by the same residue s (Cys143, Met147, His323 and Met325; Fig. 5A). The other two silver sites have lower occupancies (Ag2, ∼10 σ and Ag3, ∼10σ) and are each coordinated by two methionines of the methionine track (Met286 and Met339 for Ag2; Met341 and Met343 for Ag3). While direct measurement of the affinities of the methionine binding track sites would be challenging, the structural data suggest that the methionine track provides several low-affinity binding sites for Ag(I), and, by extension, for Cu(I). Thus, while the methionine track binds Cu(I) but not Cu(II), the high affinity CxxxM-HxM site likely binds both copper redox states.

**Figure 5.**
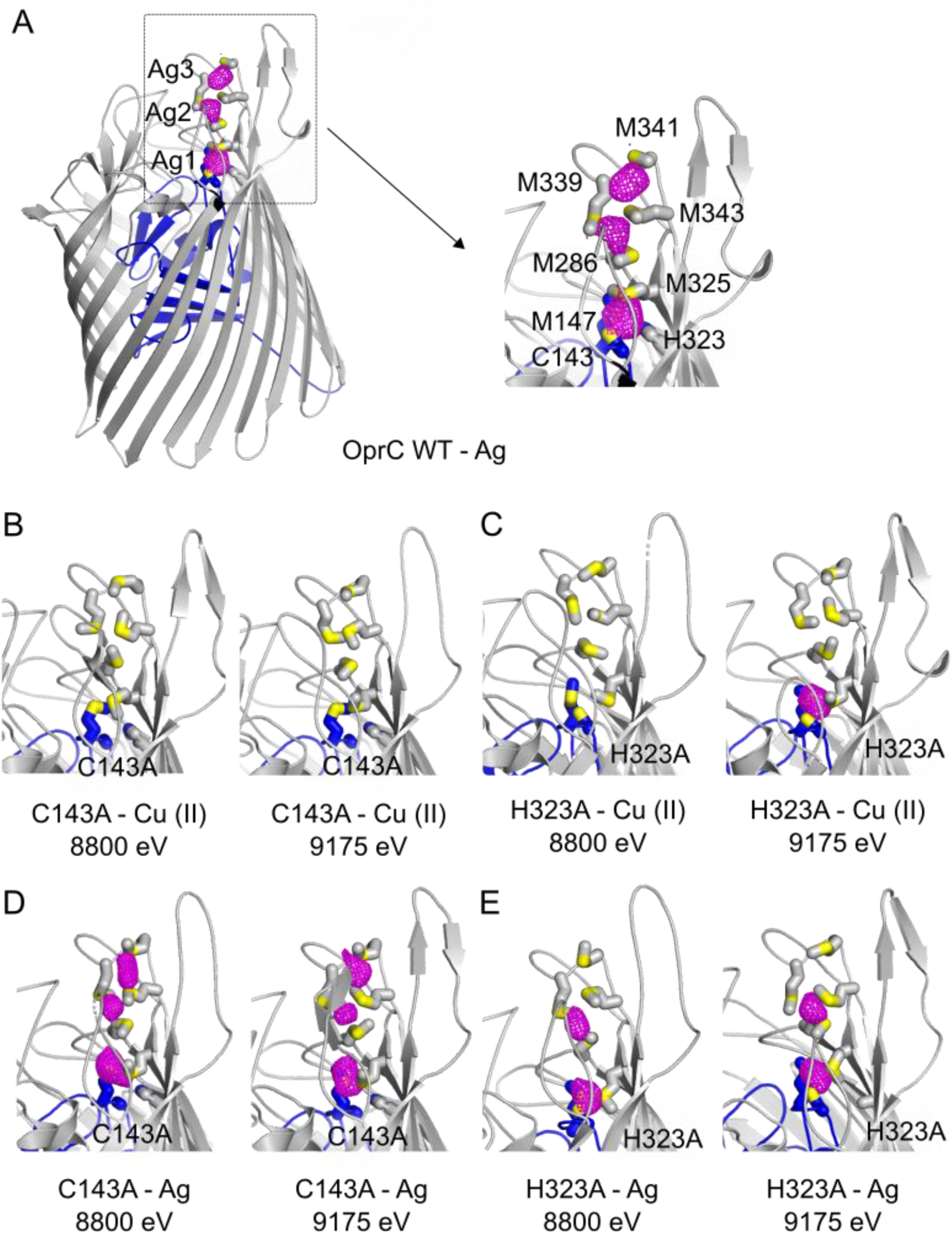
OprC binds Cu(I). (A-E) Anomalous difference maps of (A) OprC_WT_, (B,D) C143A and (C,E) H323A variants crystallised in the presence of (A,D,E) Ag or (B,C) Cu(II), and collected at different energies. The inset to (A) shows a close-up of the anomalous difference peaks (magenta) near the principal binding site in OprC_WT_, with binding residues labelled and represented as stick models. Sulphurs are coloured yellow. For clarity, the metal used in co - crystallisation and the energy used for data collection are shown underneath each panel. The OprC plug domain is coloured blue.

To obtain more information on the individual residue contribution to copper binding, we next generated the complete set of single alanine mutants of the principal binding site residues (C143A, M147A, H323A and M325A), and determined copper binding via analytical SEC and ICP-MS. For all single mutants, the copper content after purification from LB was below 10%, except for M147A (∼20 %) (Fig. 3b). Upon incubation with 3 or 10 equivalents Cu (II), various occupancies were obtained. C143A has no bound copper even after incubation with 10 equivalents Cu (II), suggesting this residue has a crucial role. H323A (∼30%) and in particular M147A (∼60%) have relatively high occupancies after copper incubation and SEC, indicating that these residues contribute less towards binding. Of the four ligands, the M147 thioether is the furthest away from the copper in the crystal structure (Fig. 2B), which may explain why it contributes the least to ligand binding. Interestingly, removal of bound copper is much faster in the M147A mutant compared to OprC_WT_ (Fig. 3C), suggesting that solvent exclusion by the M147 side chain (Fig. 4D) is the main reason why copper is kinetically trapped in OprC_WT_.

To shed additional light on the redox state of the bound copper, continuous wave EPR (cw - EPR) spectra were recorded on OprC_WT_. Surprisingly, as-purified OprC_WT_ containing ∼0.6 equivalents copper was EPR silent (Fig. 3D), demonstrating that the copper species present in the crystal structure is Cu(I). The as-purified M147A protein, with ∼0.1 equivalent copper, was EPR silent as well. We next loaded the M147A mutant with CuSO4 to 1 equivalent, and EPR spectra were recorded over time. The observed EPR signal is different from the standard CuSO4 Cu (II) EPR signals, confirming that Cu(II) binds to the protein. The EPR spectra of the OprC-WT and OprC-M147A mutant show nicely resolved ^63,65^Cu(II) hyperfine coupling along the parallel region, due to the interaction of an unpaired electron spin (*S*= ½) of Cu(II) with the nuclear spin of (*I =* 3/2*) of* ^63,65^Cu nuclei, as indicated by the blue goal-post in Fig. 3D. Interestingly, the EPR signals decrease slowly upon prolonged incubation, suggesting that bound Cu (II) is very slowly reduced to Cu(I) (Figs. 3 E,F). This, together with the possibility that OprC binds Cu(I) from the LB media, could be an explanation for the observation that as-purified OprC, expressed under aerobic conditions, contains reduced copper. However, it is clear that the observed reduction of Cu(II) is too slow to be physiologically relevant, obviating the need to find a mechanistic explanation.

### Cysteine is essential for high-affinity copper Cu (II) binding

While wild type OprC and most single alanine mutants can be (partly) loaded via Cu(II) incubation, this is not the case for the C143A mutant (Fig. 3B). Given that OprC bind s Cu(II), we hypothesised that removal of the cysteine might lead to much lower affinity for Cu(II), so that after SEC nothing remains bound. To test this we determined the crystal structures of the OprC C143A mutant co-crystallised with Cu (II) or silver Ag (I). For each crystal, datasets were collected at 8800 eV and 9175 eV. Bound copper is expected to give a strong anomalous peak only at 9175 eV (above the copper K edge at 8979 eV), while bound silver will give comparable peaks at both energies (silver L-III edge at 3351 eV). For C143A co-crystallised with Cu(II), no anomalous peaks are visible at both energies (Fig. 5B), showing that Cu (II) binding is indeed abolished. Crucially, in the presence of silver, the same three anomalous peaks are visible as for OprC_WT_ (compare Figs. 5A and D), strongly suggesting that the C143A mutant can still bind Cu(I). Since Cu(II) prefers histidine nitrogen as ligands and Cu(II) binding sites often contain one or more His residues, we also co-crystallised the H323A mutant with Cu(II) and Ag(I). As shown in Fig. 5, one strong anomalous peak, at the high -affinity binding site, is observed with Cu(II), supporting the SEC data that the histidine is not required for Cu(II) binding. With Ag(I), two clear anomalous peaks are observed, suggesting that the H323A mutant can still bind Cu(I) at the principal site and at the methionine track.

### OprC mediates copper uptake in *P. aeruginosa*

To demonstrate that OprC imports copper, we performed anaerobic growth experiments in *P. aeruginosa* with added copper. Given that *oprC* expression is repressed with excess external Cu(II) (26–29), we employed arabinose-inducible overexpression of His-tagged *oprC* via the broad range pHERD30 plasmid(35). We complemented the PA14 Δ*oprC* strain with OprC_WT_ and OprC_AA_-containing plasmids and performed growth assays in rich media with empty vector as control. Fig. S4 shows clear toxicity when OprC_WT_ is overexpressed, even without Cu(II) addition. Surprisingly, expression of OprC_AA_ was equally toxic as OprC_WT_ overexpression, which indicates that the toxicity phenotype is caused by overexpression of OprC *per se*, and is not linked to copper uptake.

Since copper toxicity assays failed, we decided to determine *P. aeruginosa* whole cell metal contents using ICP-MS. We observed no differences in copper content between the wild type PA14 and Δ*oprC* strains in rich media without added copper (Fig. S5), suggesting that OprC is not expressed under these conditions. By contrast, cells expressing OprC_WT_ from pHERD30 have more associated copper when compared to the empty vector control, under both aerobic and anaerobic conditions (Fig. 6A). However, as suggested by the toxicity phenotypes described above, this could be due to increased leakiness of cells as a result of plasmid-based OMP expression, a possibility that was not taken into account in a recent study(29). Crucial is therefore the result of cells expressing the OprC_AA_ inactive mutant, showing copper levels similar to those of the control. Moreover, OprC_WT_ and OprC_AA_ are present at similar levels in the OM (Fig. 6B), demonstrating that the different amounts of copper associated with the cells are not due to differences in protein levels. In addition, no substantial differences in cellular metal content were detected for other divalent metals. These results firmly establish OprC as a copper importer in *P. aeruginosa*.

**Figure 6.**
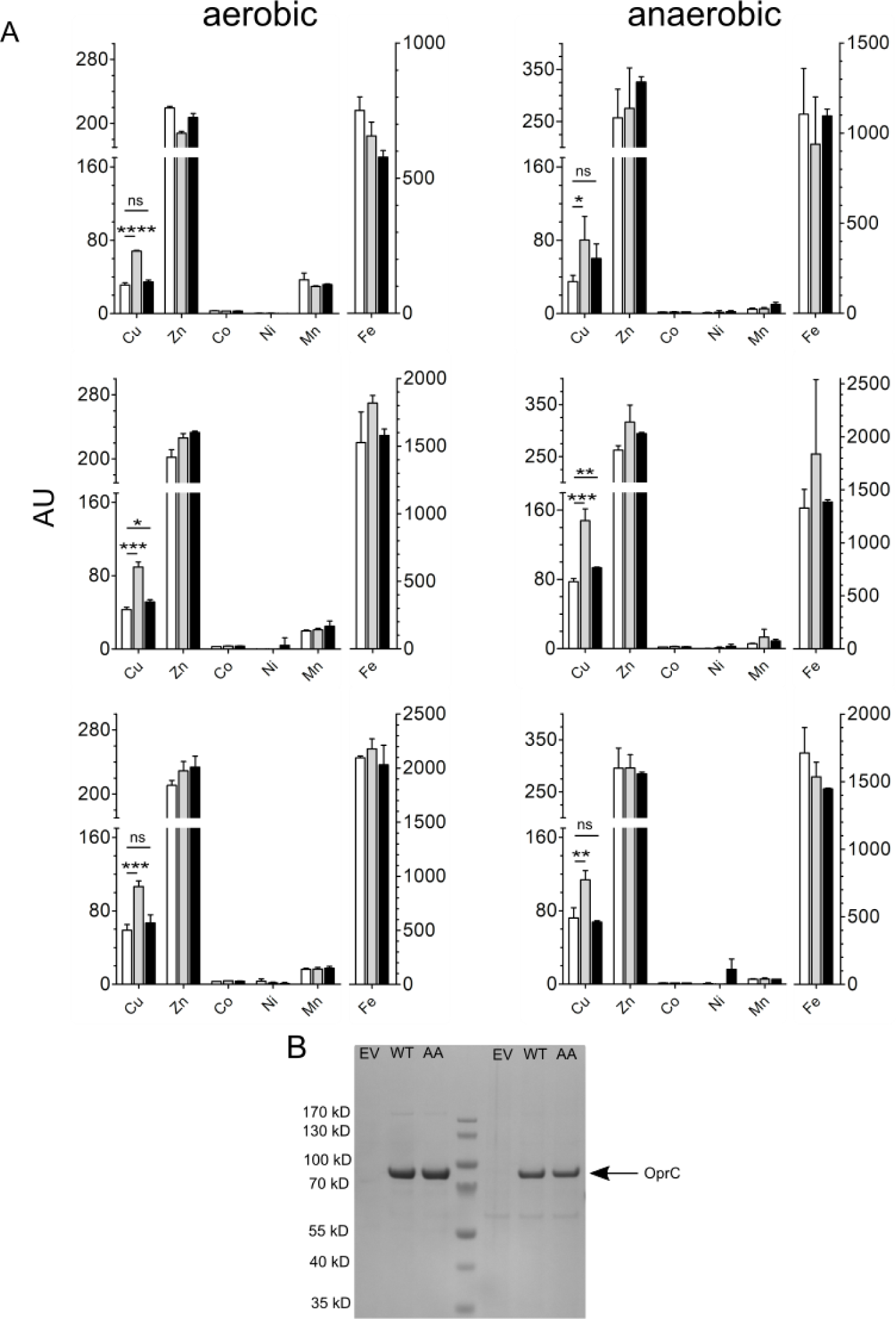
OprC is an OM copper importer. Whole-cell metal content of PA14 *ΔoprC* cells overexpressing empty vector (white bars) OprC_WT_ (grey bars) and OprC_AA_ proteins (black bars) analysed via ICP-MS. Cell associated metal content was determined in cells grown in rich media supplemented with 100 mM sodium nitrate (no added copper) under both aerobic (left panels) and anaerobic conditions (right panels). The three biological replicates are plotted separately due to differences in absolute metal levels. Reported values are averages ± s.d. (n = 3). Significant levels were analysed via unpaired two tailed t-test. ns., not significant (p ≥0.05); *, p ≤ 0.05; **, p ≤ 0.01; ***, p ≤ 0.001; ****, p ≤ 0.0001. (G) SDS-PAGE gel of pHERD30-overexpressed OprC_WT_ and OprC_AA_ proteins in PA14 *ΔoprC* after IMAC. Molecular weight marker is shown on the left.

### OprC is abundant in *P. aeruginosa* during infection

A recent study(36) suggested that OprC increases the virulence of *P. aeruginosa*, but did not provide a measure of the abundance of the transporter *in vivo*. To determine the abundance of OprC in *P. aeruginosa* UCBPP-PA14 and *Acinetobacter baumannii* ATCC 19606 in infected lung tissues, we employed a sensitive targeted proteomics approach with parallel reaction monitoring, with absolute quantification using heavy-isotope labeled reference peptides (Methods). The results showed that in both mouse and rat pneumonia models, OprC was present at 1,000 to 10,000 molecules per *P. aeruginosa* cell, making it one of the five most abundant TonB-dependent transporters. As a comparison, the most abundant TonB-dependent transporter, FpvA, had 8,000 to 33,000 molecules per cell. We also assessed OprC abundance in *A. baumannii*, due to the facts that (i) AbOprC has also been linked to virulence(37) and (ii) both proteins are highly similar (50% sequence identity, Fig. S6). Moreover, both bacteria are important human pathogens with a similar, low-permeability OM. In *A. baumannii*, OprC was less abundant in mouse and rat pneumonia models (40 to 400 molecules per cell), while the most abundant TonB-dependent transporters BfnH and BauA were present at 500 to 3’000 molecules per cell.

## Discussion

Our data show that the TBDT OprC binds copper at an unusual CxxxM-HxM site that becomes solvent excluded upon metal binding, kinetically trapping the metal and precluding determination of metal binding affinities. While our data strongly suggest that the CxxxM-HxM site can accommodate both Cu(I) and Cu(II), the observed crystal structure is that with Cu(I) bound, and the precise geometry of the binding site with Cu(II) is unknown. The copper site most similar to that of Cu(I) in OprC occurs in class I Type I copper proteins like cytochrome c oxidase, pseudoazurin and plastocyanin, electron transfer proteins that can coordinate both Cu(I) and Cu(II) via 2Cys+1Met+1His site or 2His+1Cys+1Met sites. Interestingly, the active site His117 in azurin renders the copper atom solvent inaccessible(38, 39), reminiscent to the likely role of Met147 in OprC. The superficial similarity of the OprC binding site to that of (pseudo)azurin prompted us to generate the M147H and M325H OprC mutants in an attempt to convert OprC into a unique, blue copper transport protein. However, in the presence of added Cu(II), both mutants remain colourless, and comparison of the crystal structures with those of pseudoazurin and plastocyanin show small but most likely important differences in the geometries of the active sites (Fig. S7). All four residues in the OprC binding site contribute to copper binding, but not to the same extent and, in some cases, not equally for both copper redox states. This is illustrated by Cys143, which is essential for high-affinity Cu(II) binding but dispensable for Cu(I). By contrast, removal of His323 still allows Cu(II) and Cu(I) binding. The presence of a methionine binding track, constituting several low-affinity binding sites for Ag(I)/Cu(I) but not for Cu(II), suggests that metal delivery to OprC may occur in different ways for Cu(I) and Cu(II). Methionines also coordinate Cu(I) in other copper transporters, such as the bacterial CusABC and CopA exporters and the eukaryotic Ctr1 copper importer(17, 18, 40–42). These transporters have been shown to also transport silver, but structures with silver have been determined only for the Cus system(17, 40). By analogy, our structural data therefore suggest that OprC can also import silver. The ability of OprC to acquire Cu(I) could be important for biofilms, which are anaerobic to various degrees depending on the location inside the biofilm(43, 44). Oxygen tension also reduces as lung chronic disease mediated by *P. aeruginosa* progresses, turning airway mucus into an anaerobic environment in cystic fibrosis patients that will favour the availability of Cu(I) (45). *P. aeruginosa* is capable of anaerobic respiration by using nitrate, nitrite or nitrous oxide as terminal electron acceptor (46), and OprC has been shown to be induced under anaerobic conditions(47).

With respect to Cu(II), the affinity of 2.6 µM reported by an early study (26) most likely resulted from non-specific binding, since the rich media used (48) to culture *P. aeruginosa* would have generated OprC with high copper occupancy. A recent study in *P. aeruginosa* proposed a novel copper uptake mechanism in copper-limited conditions, which involves secretion of the copper binding protein azurin by a CueR-regulated Type VI secretion system. The secreted azurin would scavenge Cu(II) from the environment and load it onto OprC via a direct interaction, conferring a competitive advantage under copper-limiting conditions(29). Delivery by azurin would be an efficient way to load OprC with Cu(II) and would presumably not require any low affinity sites to direct the metal to the principal binding site as for Cu(I). However, the pulldown experiment done by Han et al. to show the azurin-OprC interaction was done with OprC folded *in vitro* from inclusion bodies (29). Given that TBDTs are hard to fold *in vitro* due to their large size and complex architecture, and no attempts were made to assess the functionality of the obtained OprC, its interaction with azurin remains to be confirmed. In our hands, no complex formation between OM-purified, functional OprC and azurin was observed via SEC, suggesting any interaction will be transient.

Given that copper, and in particular the more toxic Cu(I), is a known antimicrobial, the presence of bacterial proteins dedicated to copper acquisition such as OprC might be problematic under certain conditions. Indeed, it is thought that, in contrast to iron that is withheld from a pathogen by the host during infection, elevated levels of host-derived copper in *e.g*. macrophages could be an alternative “nutritional immunity” antimicrobial response (49). In this model, bacterial virulence would be attenuated by mutations, particularly in transporters, that cause copper sensitivity(49). However, recent data suggest that deletion of *oprC* results in reduced quorum sensing, impaired motility and lower virulence of *P. aeruginosa*, leading to the proposal that the presence of OprC is critical for virulence(36). In addition, another recent study reported decreased virulence of an *A. baumannii oprC* knockout(37). The decreased virulence of *oprC* knockouts in these studies appears at odds with what one would expect from the copper nutritional immunity model(49), as is our proteomics data showing that OprC is very abundant in a *P. aeruginosa* mouse infection model. While we did not assess *oprC*-dependent virulence, these data do suggest that copper is withheld by the host under many experimental conditions. While much still needs to be learned, it is clear that regulation of any copper import protein is crucial, possibly both at the gene and protein level. Unfortunately, and in contrast to the many copper stress genes that, as part of the CopR or CueR regulons, are upregulated under aerobic conditions during copper stress(9, 11, 27, 28, 50, 51), nothing is known about how *oprC* is downregulated during such stress. Intriguingly, *oprC* (*PA3790*) is in an operon with *PA3789*, which encodes for an uncharacterised inner membrane protein containing PepSY domains, hinting at a peptidase function(27). Another protein strongly downregulated during copper stress is PA5030, which is an MFS transporter with a large number of His+Met residues (26 out of 438 residues), suggesting it could mediate copper delivery to the cytoplasm, possibly in concert with OprC and an as yet unidentified periplasmic protein(27).

OprC is the first example of a TBDT that mediates copper import without a metallophore. The TBDT with the closest substrate specificity to OprC is the ionic zinc transporter ZnuD from *Neisseria meningitidis*, the structure of which has been solved(52). Large structural differences between OprC and ZnuD exist for the extracellular loops (overall Cα r.m.s.d. ∼5.9 Å). ZnuD has several discrete low-affinity binding sites that may guide the metal towards the high-affinity binding site (52). In OprC, a distinctive “methionine track” provides low-affinity binding sites to guide copper to the high affinity site. Interestingly, while the extracellular loops between OprC and ZnuD are very different and the overall sequence identity is only 28%, the metal binding sites are located at very similar positions and only 2.8 Å apart (Fig . S8), suggesting that the transport channel formed via TonB interaction may be similar. Inspection of the ZnuD structure shows that the zinc binding site is excluded from solvent, and we propose that the zinc ion in ZnuD is kinetically trapped, analogous to copper in OprC.

OprC shares ∼60 % identity to NosA from *Pseudomonas stutzeri*, for which no structure is available. Like OprC, NosA is expressed under anaerobic conditions and repressed in the presence of µM concentrations Cu(II) (53–55). *P. stutzeri* NosA antibodies did not react with *P. aeruginosa* (53), but our structure identifies NosA as an OprC ortholog, since the CxxxM-HxM copper binding motif and some of the methionine track residues are conserved (Fig. S6). NosA is important during denitrification in *P. stutzeri* JM300 and was proposed to load copper either directly or indirectly to the periplasmic N2O reductase NosZ (53–55). However, a more recent report for a different *P. stutzeri* strain found no difference between NosZ activity and copper content for a *nosA* knockout (56). In addition, OprC/NosA also occurs in a number of non-denitrifying *Proteobacteria* such as *Salmonella enterica, Klebsiella pneumoniae* and *Acinetobacter baumannii* (Fig. S6), showing that NosZ maturation is not a general function of OprC. The occurrence of OprC in some (*e.g*. *S. enterica*) but not in other (*e.g. E. coli*) Enterobacteria is intriguing, given that the OM of all Enterobacteria is relatively permeable to small molecules due to abundant general porins such as OmpC (57).

## Methods

### Recombinant production of *Pseudomonas aeruginosa* OprC

The mature version of the gene coding for *oprC* of *P. aeruginosa* PAO1 (UniProt ID; PA3790)(58), starting with His56 as determined by Nakae et al.(26), was synthesized to include a 7 x His tag at the N-terminus (Eurofins, UK), cloned into the pB22 arabinose-inducible expression vector(59) and transformed into chemically competent *Escherichia coli* DH5α cells. After expression and processing by signal peptidase, the N-terminal sequence of this construct is NVRLQHHHHHHHLEAEEHSQHQ-. A second version of this construct was constructed in a pB22 version containing a tobacco etch virus (TEV) site after the His7-tag. Correct sequences were confirmed by DNA sequencing (Eurofins, UK) using both forward and reverse plasmid - specific primers. The OprC_AA_ mutant was produced by changing the key amino acids Cys143 and Met147 to alanine residues using the KLD Quickchange site-directed mutagenesis kit (New England Biolabs, UK) and specific primers containing both mutation sites (forward: 5’ - tcgcgcggatgcaccaaccagctatattagc-3’; reverse: 5’-ttcggggcggcgccaagcatcatgc-3’). The single mutants C143A, M147A, H323A, M325A, M147H and M325H were made in similar ways.

OprC recombinant protein production and purification was performed as follows: *E. coli* C43 Δ*cyo* was electroporated with expression vector, recovered for 60 minutes in LB (Sigma, UK) at 37 °C, and plated on LB agar (Sigma, UK) containing 100 µg mL^-1^ ampicillin (Melford, UK). Transformants were cultured in LB medium or in LeMasters-Richards (LR) minimal medium with glycerol (2-3 g/l) as carbon source. All media contained 100 µg mL^-1^ ampicillin. For rich media, cells were grown (37 °C, 180rpm) until OD600 ∼0.6, when protein expression was induced with 0.1% arabinose for 4-5 h at 30 °C or overnight at 16 °C (150 rpm). For LR media, a small overnight pre-culture in LB was used at 1/100 v/v to inoculate an LR-medium pre-culture early in the morning (typically 1 ml preculture for 100 ml of cells was used), which was grown during the day at 37 °C. After late afternoon inoculation, large-scale cultures (typically 6-8 l) were grown overnight at 30 °C until OD 0.4-0.7, followed by induction with 0.1% arabinose at 30 °C for 6-8 hours. Cells were harvested by centrifugation (5,000 rpm, 20 minutes, 4 °C), and pellets homogenized in 20 mM Tris (Sigma), 300 mM NaCl (Fisher) pH 8.00 (TBS buffer), in the presence of 10 mM ethylenediamine tetra-acetic acid (EDTA, Sigma). Cells were broken by one pass through a cell disruptor (Constant Systems 0.75 kW operated at 23 kpsi), centrifuged at 42,000 rpm for 45 minutes at 4 °C (45Ti rotor; Beckman), and the resulting total membrane fraction was homogenized in TBS buffer containing 1.5% Lauryl-dimethylamine oxide (LDAO) (Sigma, UK). Membrane proteins were extracted by stirring (60 minutes, 4 °C), centrifuged (42,000 rpm in 45Ti rotor, 30 minutes, 4°C), and the membrane extract was loaded on a Chelating Sepharose Fast Flow bed resin (∼10 ml; GE Healthcare, UK) previously activated with 200 mM NiCl2 (Sigma) and equilibrated in TBS containing 0.15 % n-dodecyl-beta-D-maltopyranoside (DDM). After washing with 15 column volumes buffer with 30 mM imidazole, protein was eluted with 0.25 M imidazole buffer (Fisher), incubated with 20 mM EDTA (30 minutes, 4 °C), and loaded on a Superdex 200 16/600 size exclusion column equilibrated with 10 mM Hepes, 100 mM NaCl, 0.05% DDM, 10 mM EDTA, pH 7.5. Peak fractions were pooled and concentrated using a 50 MWCO Amicon filter (Millipore, UK), analyzed on SDS-PAGE, flash-frozen in liquid nitrogen and stored at -80C. Typical yields of purified wild type and most mutant OprC proteins ranged between 2-5 mg per l media grown at 16 °C. All media and buffer components were made in fresh milli-Q water.

Protein preparations intended for crystal trials were pooled and buffer-exchanged to 10 mM Hepes 100 mM NaCl, 0.4% tetraethylene glycol mono-octyl ether (C8E4) (Anatrace, US), pH 7.5. NaNO3 was substituted for NaCl for protein preparations intended for crystal trials with silver in order to avoid formation of insoluble AgCl. Protein preparations to be used for metal analysis after removal of the His-tag underwent a slightly different protocol. The elution fraction from immobilized metal affinity chromatography (IMAC) was buffer-exchanged to 50 mM Tris, 0.5 mM EDTA, 0.2 mM TCEP, 100 mM NaCl, 0.05% DDM (Anatrace, US), pH 7.50, and submitted to TEV protease digestion (ratio 1 mg TEV: 10 mg protein, 4 °C, overnight). Samples were submitted to a second IMAC column, where flow-through and wash fractions were combined for the subsequent SEC step in 10 mM HEPES 100 mM NaCl 0.05 % DDM 0.5 mM EDTA pH 7.5. Protein concentration was determined by BCA assay (Thermo Scientific, UK) and by UV/Vis absorbance at 280nm (considering OprC E_0.1%_ = 1.6 as determined by ProtParam).

### In vitro metal binding assays and Inductively Coupled Plasma Mass Spectrometry (ICP-MS)

OprC samples intended for metal binding assays were exchanged into respective chelex-treated buffers without EDTA and were equilibrated with different equivalents of Cu(II) or Zn(II), for 30 min at room temperature (n=3). Protein concentrations used were in the range of 10 – 20 µM. Samples were loaded on an analytical Superdex 200 Increase 10/300GL (GE Healthcare) column, equilibrated in 10 mM Hepes, 100 mM NaCl, 0.05% DDM, 0.5 mM EDTA pH 7.5. Size exclusion peaks were pooled, concentrated and quantified for protein by UV absorbance at 280 nm. Protein samples were diluted 10-fold in 2.5% HNO3. Analytical metal standards of 0 – 500 ppb were prepared by serial dilution from individual metal stocks (VWR, UK) and were matrix-matched to protein samples. Samples and standard curves were analysed by inductively coupled plasma mass spectrometry (ICP-MS) using Durham University Bio-ICP-MS Facility (Thermo X-series instrument, Thermo Fisher Scientific; PlasmaLab Software) running in standard mode (for ^55^Mn, ^59^Co, ^60^Ni, ^65^Cu, ^66^Zn) or collision cell mode (for ^56^Fe). OprC WT samples were screened for the presence of ^56^Fe, ^55^Mn, ^59^Co,

^60^Ni, ^65^Cu, ^66^Zn and rest were typically screened for the presence of ^65^Cu and ^66^Zn. The increase in copper content in “as purified” tagless WT protein (Fig. 3b) is most likely due to the use of a second IMAC column after tag cleavage. In addition to the additional handling steps that could have increased copper content, the NiCl2 used for the IMAC column might contain traces of copper.

### Copper extraction (demetallation) experiments

OprC_WT_, OprC_AA_ and M147A samples were incubated with 3 equivalents copper for 30 min and then loaded onto a Superdex S-200

Increase 10/300GL column equilibrated with 10 mM HEPES 100 mM NaCl 0.05 % DDM 0.5 mM EDTA pH 7.5. Peak fractions were pooled, concentrated and quantified by UV absorbance at 280 nm. Samples were exchanged into respective chelex-treated buffer without EDTA. For demetallation experiments, 20 µM of copper-bound proteins were taken in duplicates and incubated with 100-fold excess of the copper chelator bathocuproine disulfonate (BCS) (Sigma) and 100-fold excess of the reducing agent hydroxyl amine (NH2OH) (Sigma) at 60 °C and room temperature. BCS is a high-affinity Cu(I) chelator (logβ2 20.8) and forms a 2:1 complex with Cu(I), namely [Cu(BCS)2]^-3^, with a molar extinction coefficient of 13,300 cm^-1^ M^-1^ at 483nm, enabling quantitation of Cu(I)(60).

### Protein crystallization, data collection and structure determination

Sitting-drop crystallization trials were set up using a Mosquito crystallization robot (TTP Labtech) with commercial screens (MemGold1 and MemGold2, Molecular Dimensions) at 20 °C. To obtain the initial structure of Cu-bound OprC, the protein (∼12 mg/ml) was incubated with 3 mM CuCl_2_ for 1 hr at room temperature, followed by setting up crystallisation trials. A number of initial hits were obtained and were subsequently optimised by manual hanging drop vapour diffusion using larger drops (typically 1-1.5 ul protein + 1 ul reservoir). Well-diffracting crystals (∼3 Å resolution at a home source) were obtained in 0.1 M NaCl/0.15 M NH4SO4/0.1 M MES pH 6.5/18-22% PEG1000. Crystals were cryoprotected with mother liquor lacking CuCl2 containing 10% PEG400 for ∼5-10 s and flash-frozen in liquid nitrogen. Diffraction data were collected at Diamond Light Source (Didcot, UK) at beamline i02. For the best crystal, belonging to space group C2221, 720 degrees of data were collected at an energy of 8994 eV, corresponding to the K-edge of copper (Table S1). Data were autoprocessed by xia2(61) . The structure was solved via single anomalous dispersion (SAD) via AUTOSOL in Phenix(62). Two copper sites were found, one for each OprC molecule in the asymmetric unit (Fig. S1). The phases were of sufficient quality to allow automated model building via Phenix AUTOBUILD, generating ∼60% of the structure and using data to 2.0 Å. The remainder of the structure was built manually, via iterative cycles of refinement in Phenix and model building in COOT(63). Metal coordination was analysed by the Check-my-metal server(64). The final refinement statistics are listed in Table S1. Subsequently, crystals were also obtained without any copper supplementation of the protein. These were isomorphous to those described above and obtained under identical conditions. Molecular replacement indicated the presence of copper and an identical structure to that obtained above (data not shown). OprC_AA_ crystals (∼10 mg/ml protein) were obtained and optimized by hanging drop vapor diffusion as described above, and diffraction-quality crystals were obtained in the same conditions as for Cu-OprC, *i.e*. 0.1 M sodium chloride/0.15 M ammonium sulfate/0.01 M MES sodium pH 6.5/19% (w/v) PEG1000. Interestingly however, the OprC_AA_ crystals belong to a differentspace group (P22121), most likely as a result of the structural differences between both OprC variants. Diffraction data were collected at Diamond Light Source (Didcot, UK) at beamline I24. Diffraction data were processed in XDS (65). The structure was solved by molecular replacement (MR) using Phaser, with wildtype OprC as the search model. Model building wa s done in COOT and refinement in Phenix. As for Cu-OprC, the data collection and refinement statistics are shown in Table S1. C143A and H323A proteins (∼10-12 mg/ml protein) were incubated with 2 mM CuSO4 at room temperature for 1 hr, followed by co-crystallisation. Diffracting crystals for both C143A and H323A in the presence of copper were obtained in 0.34 M Ammonium sulfate/0.1 M Sodium citrate pH 5.5/12 -16 % w/v PEG 4000 and were cryo-protected using mother liquor lacking CuSO4 and with 25% ethylene glycol for ∼10 s and flash-frozen in liquid nitrogen. M147H and M325H crystals were obtained in the same condition as those for wild type OprC. For co-crystallisation with silver, OprC proteins were incubated with 2 mM AgNO3 for 1 hour at room temperature, followed by co-crystallisation. Well-diffracting OprC_WT_ crystals with silver were obtained under the same conditions as in the presence of copper. For the best OprC_WT_ crystal, belonging to space group C2221, 999 degrees of data were collected at an energy of 8000 eV to obtain anomalous signals for Ag. C143A and H323A crystals (∼10-12 mg/ml protein) with Ag were obtained from 0.2 M Choline chloride/0.1 M Tris pH 7.5/12-16 % w/v PEG 2000 MME and 0.5 M Potassium chloride/0.05 M HEPES pH 6.5/12-16 % v/v PEG 400, respectively. Crystals were cryoprotected for 5-20 s with mother liquor lacking AgNO3 but containing 25% ethylene glycol for C143A and 20 % PEG 400 for H323A. For C143A and H323A crystallised in the presence of copper or silver, datasets of 360 degrees each were collected at energies of 8800 and 9175 eV, using different parts of the same crystal (Tables S2 and S3).

### Electron Paramagnetic Resonance Spectroscopy

Electron Paramagnetic Resonance (EPR) measurements were carried out using a Bruker ELEXSYS-E500 X-band EPR spectrometer operating in continuous wave mode, equipped with an Oxford variable-temperature unit and ESR900 cryostat with Super High-Q resonator. All EPR samples were prepared in quartz capillary tubes (outer diameter; 4.0 mm, inner diameter 3.0 mm) and frozen immediately in liquid N2 until further analysis. The experimental setup and conditions were similar to those reported previously(66). The low temperature EPR spectra were acquired using the following conditions: sweep time of 84 s, microwave power of 0.2 mW, time constant of 81 ms, average microwave frequency of 9.44 GHz and modulation amplitude of 5 G, T = 20 K. The concentration of OprC_WT_ and M147A varied from 210-260 μM in 10 mM HEPES, 100 mM NaCl, 0.03 % DDM (n-dodecyl-D-maltoside), pH 7.5.

### Determination of whole cell metal content

Whole cell metal content was determined as described previously. Briefly, overnight bacterial cultures of overexpressed OprC WT and OprC AA (with empty vector as control) in PA14 *ΔoprC* background were diluted with 1:100 fresh LB supplemented with 100 mM NaNO3, and were grown to an OD of around 1.0 at 37 C. 25 ml cultures were pelleted and were washed twice in TBS and once in 20 mM Tris 0.5 M sorbitol and 200 uM EDTA pH 7.5. The cell pellets were digested in 1 ml of 68 % conc. nitric acid for > 24h. Digested sample pellets were diluted 10 fold in 2 % nitric acid (prepared in chelex-treated milli-Q water) and were analyzed by ICP-MS. Results were corrected for ODs and dilution factors. Protein levels in the OM were verified by IMAC. Briefly, 0.5 liter of OprC_WT_ and OprC_AA_ overexpressing strains (with empty vector as control) in the *P. aeruginosa* Δ*oprC* background strain were grown in LB (supplemented with 100 mM NaNO3 and 0.1% arabinose) for 6h to OD600 ∼1.0, followed by cell harvesting, cell lysis and purification as described above for *E. coli*.

### *In vivo* metal toxicity assays

For metal toxicity assays in *P. aeruginosa*, overexpressed OprC WT, C143A and AA strains using broad range plasmid pHERD30 (with empty vector as control) in PA14 *ΔoprC* were used and assays were performed in anaerobic conditions. The Cu(II) (CuSO4) range tested varied from 0-7 mM. Cultures in triplicates were inoculated with 1:100 of the pre-cultures grown in anaerobic conditions (LB with 100 mM sodium nitrate). Growth curves of final volume 200 µl were set up in 96-well Costar culture plate (Sigma Aldrich) and sealed inside an anaerobic chamber (Don Whitley Scientific, A35 workstation). Growth was monitored at 600 nm using an Epoch plate reader (Biotek Instruments Ltd) at 37 °C. Time points were collected with 30 min intervals and experiments were performed in triplicates.

### Animal infection models

*Intra-tracheal instillation model*: specific pathogen free (SPF) immunocompetent male Sprague-Dawley rats weighing 100 - 120 g or male CD-1 mice weighing 20 - 25 g were infected by depositing an agar bead containing around 10^7^ colony-forming units *Acinetobacter baumannii* ATCC 19606 and *Pseudomonas aeruginosa* UCBPP-PA14, deep into the lung via nonsurgical intra-tracheal intubation(67). In brief, animals were anesthetized with isoflurane (5%) and oxygen (1.5 L/min) utilizing an anesthesia machine. Depth of anesthesia was evaluated by absence of gag reflex; if the reflex was present, the animal was placed back under anesthesia until the reflex disappeared. No animals were utilized until they were fully anesthetized. Animals were infected via intra-bronchial instillation of molten agar suspension (rats-100 µl) (mice-20 µl) via intra-tracheal intubation, and then allowed to recover. Animals were returned in their home cages and observed until recovered from anesthesia. At 24 h post infection, animals were sacrificed and lung was homogenized in sterile saline using a lab blender. All procedures are in accordance with protocols approved by the GSK Institutional Animal Care and Use Committee (IACUC), and meet or exceed the standards of the American Association for the Accreditation of Laboratory Animal Care (AAALAC), the United States Department of Health and Human Services and all local and federal animal welfare laws.

### Sample workup for proteomics

The sample workup protocol was optimized to deplete host material while maintaining *A. baumannii* and *P. aeruginosa* viability until lysis. All buffers and equipment were used at 0 to 4 °C to minimize proteome changes during sample workup. The sample volume (maximum of 1 ml) was estimated and an equal volume of 1% Tergitol in PBS was added followed by vigorous vortexing for 30 s. After centrifugation at 500 x g for 5 min, the supernatant was transferred to a fresh tube, and the pellet was extracted again with 2 ml 0.5% Tergitol in PBS.

The supernatant was combined with the first supernatant and centrifuged at 18’000 x g for 5 min. The pellet was washed with 2 ml and again centrifuged at 18’000 x g for 5 min. The supernatant was removed, and the pellet was resuspended in 100 µL 5% sodium deoxycholate, 5 mM Tris (2-carboxyethyl) phosphine hydrochloride, 100 mM NH4HCO3. The sample was incubated at 90°C for 1 min. and then stored at -80 °C. Samples were thawed and sonicated for 2 x 20 s (1 s interval, 100% power). Proteins were alkylated with 10 mM iodoacetamide for 30 min in the dark at room temperature. Samples were diluted with 0.1M ammonium bicarbonate solution to a final concentration of 1% sodium deoxycholate before digestion with trypsin (Promega) at 37°C overnight (protein to trypsin ratio: 50:1). After digestion, the samples were supplemented with TFA to a final concentration of 0.5% and HCl to a final concentration of 50 mM. Precipitated sodium deoxycholate was removed by centrifugation at 4°C and 14’000 rpm for 15 min. Peptides in the supernatant were desalted on C18 reversed phase spin columns according to the manufacturer’s instructions (Macrospin, Harvard Apparatus), dried under vacuum, and stored at −80°C until further processing.

### Parallel reaction monitoring

Heavy proteotypic peptides (JPT Peptide Technologies GmbH) were chemically synthesized for *A. baumannii* and *P. aeruginosa* outer membrane proteins. Peptides were chosen dependent on their highest detection probability and their length ranged between 7 and 20 amino acids. Heavy proteotypic peptides were spiked into each sample as reference peptides at a concentration of 20 fmol of heavy reference peptides per 1 µg of total endogenous protein mass. For spectrum library generation, we generated parallel reaction -monitoring (PRM)(68) assays from a mixture containing 500 fmol of each reference peptide. The setup of the μRPLC-MS system was as described previously(69). Chromatographic separation of peptides was carried out using an EASY nano-LC 1000 system (Thermo Fisher Scientific) equipped with a heated RP-HPLC column (75 μm x 37 cm) packed in-house with 1.9 μm C18 resin (Reprosil-AQ Pur, Dr. Maisch). Peptides were separated using a linear gradient ranging from 97% solvent A (0.15% formic acid, 2% acetonitrile) and 3% solvent B (98% acetonitrile, 2% water, 0.15% formic acid) to 30% solvent B over 60 minutes at a flow rate of 200 nl/min. Mass spectrometry analysis was performed on Q-Exactive HF mass spectrometer equipped with a nanoelectrospray ion source (both Thermo Fisher Scientific). Each MS1 scan was followed by high-collision-dissociation (HCD) of the 10 most abundant precursor ions with dynamic exclusion for 20 seconds. Total cycle time was approximately 1 s. For MS1, 3e6 ions were accumulated in the Orbitrap cell over a maximum time of 100 ms and scanned at a resolution of 120,000 FWHM (at 200 m/z). MS2 scans were acquired at a target setting of 1e5 ions, accumulation time of 50 ms and a resolution of 30,000 FWHM (at 200 m/z). Singly charged ions and ions with unassigned charge state were excluded from triggering MS2 events. The normalized collision energy was set to 35%, the mass isolation window was set to 1.1 m/z and one microscan was acquired for each spectrum.

The acquired raw-files were converted to the mascot generic file (mgf) format using the msconvert tool (part of ProteoWizard, version 3.0.4624 (2013-6-3)). Converted files (mgf format) were searched by MASCOT (Matrix Sciences) against normal and reverse sequences (target decoy strategy) of the UniProt database of *Acinetobacter baumannii* strains ATCC 19606 and ATCC 17978, and Pseudomonas aeruginosa UCBPP-PA14, as well as commonly observed contaminants. The precursor ion tolerance was set to 20 ppm and fragment ion tolerance was set to 0.02 Da. Full tryptic specificity was required (cleavage after lysine or arginine residues unless followed by proline), three missed cleavages were allowed, carbamidomethylation of cysteins (+57 Da) was set as fixed modification and arginine (+10 Da), lysine (+8 Da) and oxidation of methionine (+16 Da) were set as variable modifications. For quantitative PRM experiments the resolution of the orbitrap was set to 30,000 FWHM (at 200 m/z) and the fill time was set to 50 ms to reach a target value of 1e6 ions. Ion isolation window was set to 0.7 Th (isolation width) and the first mass was fixed to 100 Th. Each condition was analyzed in biological triplicates. All raw-files were imported into Spectrodive (Biognosys AG) for protein and peptide quantification.

## Author contributions

BvdB designed the study. SPB and BvdB expressed, purified, and crystallized proteins. AB collected the diffraction data. SPB and BvdB analysed the diffraction data and refined the structures. SPB performed metal binding and *in vivo* growth experiments. TRY performed ICP-MS analyses. MS performed cw-EPR measurements and interpreted the data. SH performed proteomics experiments, supervised by DB. BvdB and SPB wrote the paper.

## Acknowledgements

SPB is supported by a Biotechnology and Biological Sciences Research Council (BBSRC, UK) grant (BB/R004366/1 to BvdB). We would like to acknowledge Scott Sucoloski, Jennifer Hoover, Josh West (Glaxo Smith Kline) for providing proteomics samples. We thank Bastien Belzunces, Chris Skylaris and Syma Khalid (University of Southampton) for exploratory quantum chemical calculations. We also thank Kevin Waldron (Newcastle University) for useful discussions and for carrying out initial ICP-MS analyses. We also thank Deenah Osman and Nigel Robinson (Durham University) for ICP-MS analyses and helpful discussions, supported by awards BB/L009226/1 and BB/R002118/1 from the BBSRC. TRY was supported by a Research Fellowship from the Royal Commission for the Enhibition of 1851. We are indebted to the Diamond Light Source for beam time (proposals mx9948, mx13587 and mx18598) and beamline assistance. MS acknowledges the EPSRC National (UK) EPR Research Facility and Service for use of the EPR spectrometers. MS and BvdB thank Luisa Ciano for the useful discussions at an early stage. The research leading to these results was in part conducted as part of the Translocation consortium (www.translocation.eu) and has received support from the Innovative Medicines Initiatives Joint Undertaking under Grant Agreement No. 115525, resources that are composed of financial contributions from the European Union’s seventh framework programme (FP7/2007 –2013) and European Federation of Pharmaceutical Industries and Associations companies in-kind contribution. BvdB would also like to acknowledge the Royal Society for salary support.

## Accession codes

Coordinates and structure factors have been deposited in the Protein Data Bank (http://www.ebi.ac.uk/pdbe/) with accession codes 6FOK (OprC_WT_), 6FOM (OprC_AA_), 6Z8Q (OprC_WT_ Ag 8000 ev), 6Z9I (OprCC143A Ag 8800 eV), 6Z99 (OprCC143A Ag 9175 eV), 6Z8Y (OprCC143A Cu 8800 eV), 6Z8Z (OprCC143A Cu 9175 eV), 6Z8T (OprCH323A Ag 8800 eV), 6Z8U (OprCH323A Ag 9175 eV), 6Z8R (OprCM147H), 6Z8S (OprCM325H), 6Z9N (OprCH323A Cu 9175 eV), 6Z9Y (OprCH323A Cu 8800 eV).

## Supporting Information

**Figure S1.**
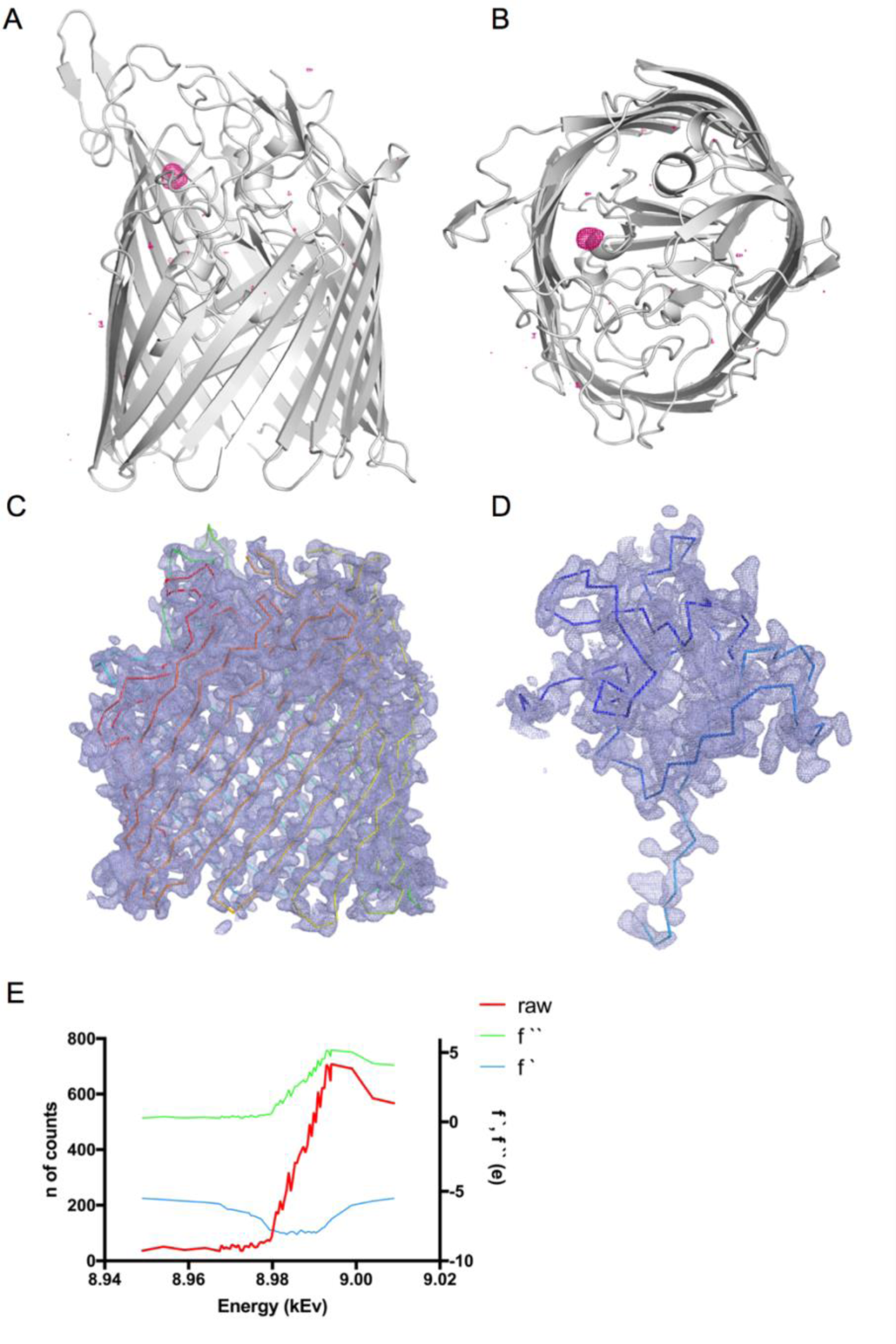
Anomalous data for the OprC_WT_ Cu-SAD experiment. (A, B) Copper anomalous maps (coloured magenta) contoured at 4 σ (carve = 30). Experimental density for one OprC protomer after density modification (but before model building) for (C) barrel and (D) N -terminal plug domain (map contoured at 1.5 σ, carve = 2.0). Ribbon is shown for orientation purposes. (E) X-ray fluorescence spectrum showing the copper-specific energy peak.

**Figure S2.**
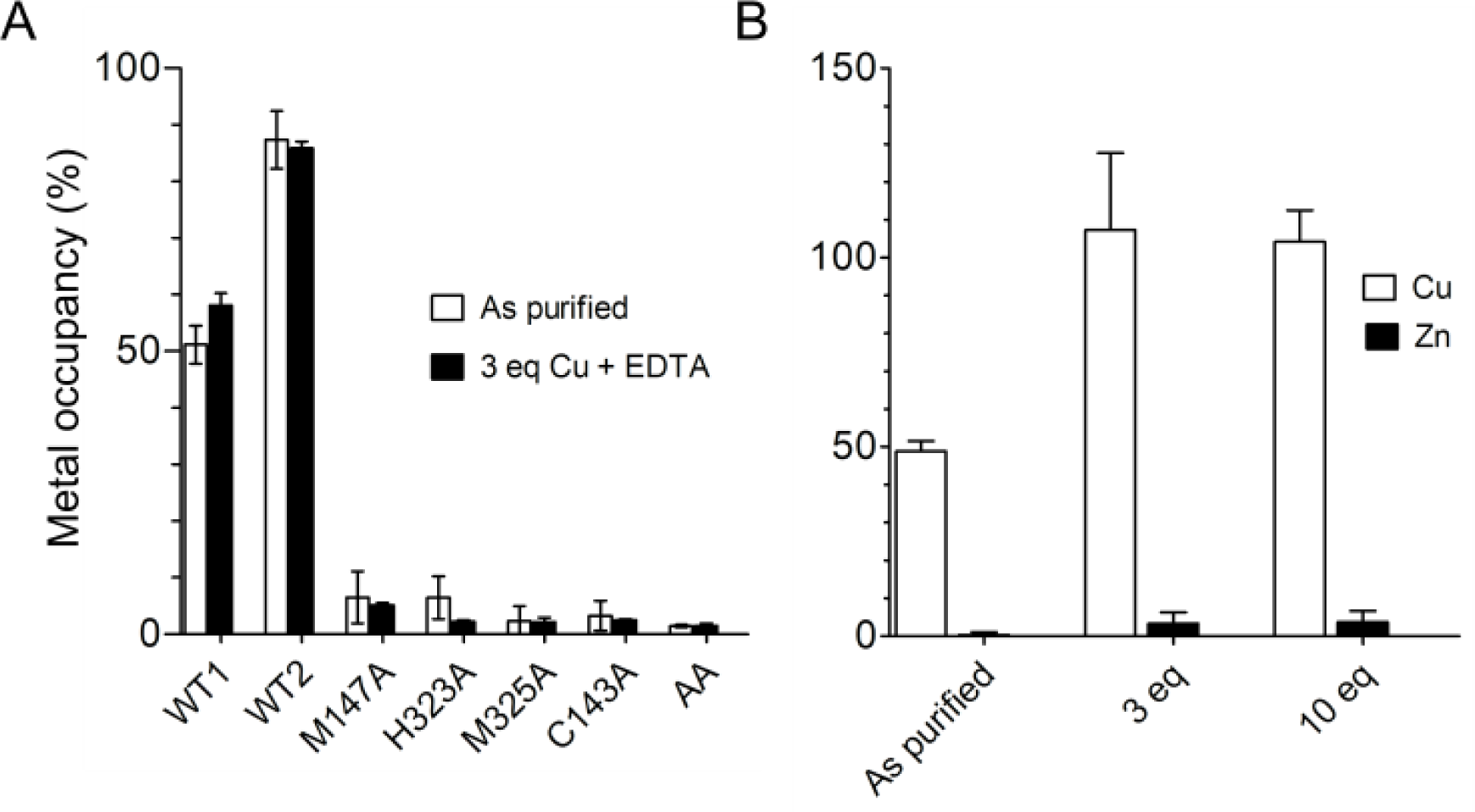
ICP-MS data of OprC and mutant proteins. (A) Metal occupancy of OprC and mutant proteins after incubation with 3 equivalents copper in the presence of 0.5 mM EDTA (∼50-fold excess) followed by analytical size exclusion chromatography and subsequent metal analysis by ICP-MS. (B) Metal occupancy of OprC WT after incubation with 3 or 10 Eq. of Cu or Zn.

**Figure S3.**
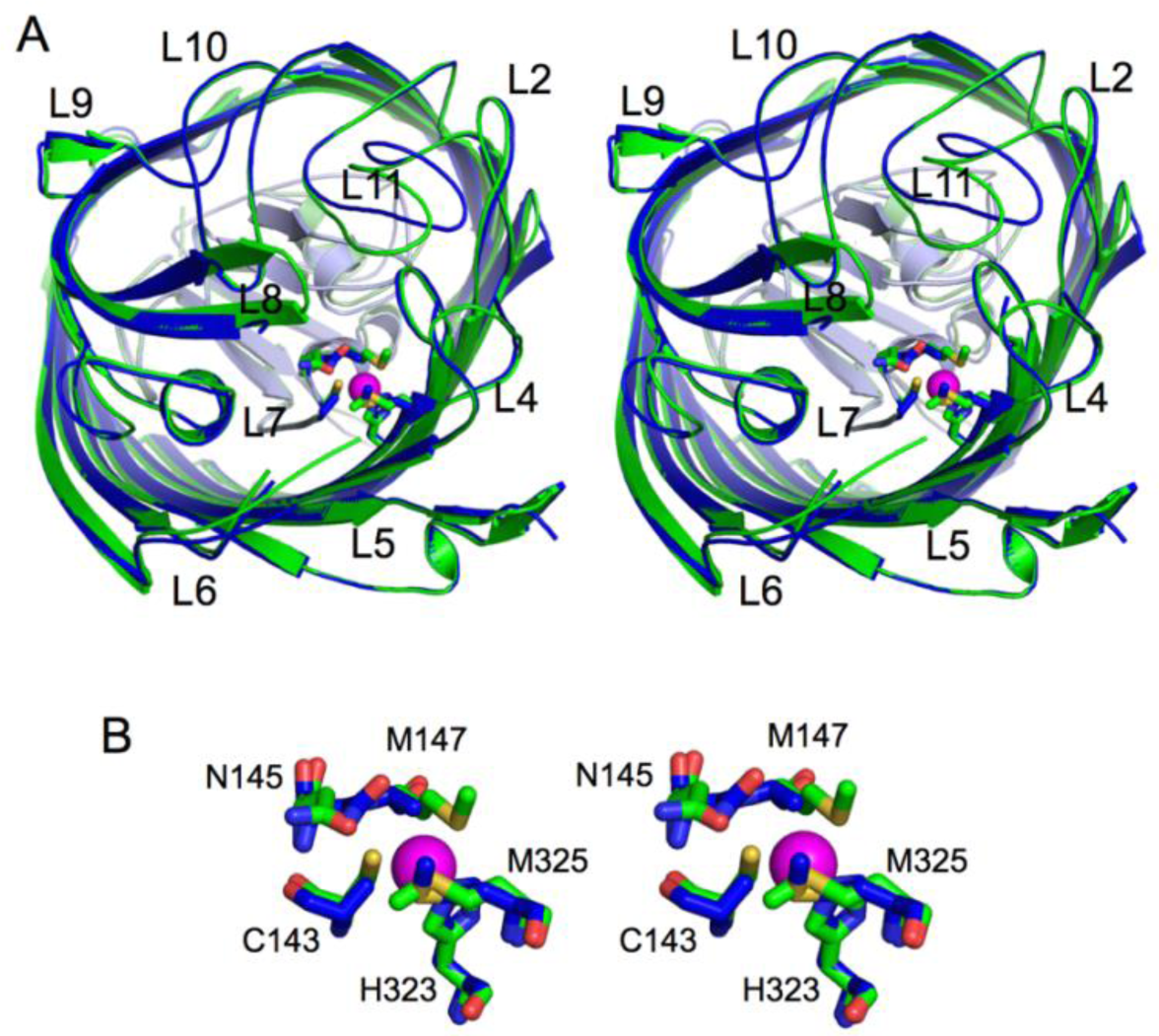
Stereo 3D representation of superposed OprC (coloured green) and OprC_AA_ (blue). (A) Extracellular view showing loops 2, 4, 5, 6, 7, 8, 9, 10, 11 (L2, L4, L5, L6, L7, L8, L9, L10, L11). Conformational changes are observed for external loops L8 and L11. (B) Active site view illustrating superposed residues involved in metal coordination for wild type OprC (green sticks) and OprC_AA_ (blue sticks). Asn145 (N145) is also shown due to its role in shielding the active site. Oxygen atoms in amino acid residues are coloured red, nitrogens blue and sulphurs yellow. Copper atom is represented as a magenta sphere.

**Figure S4.**
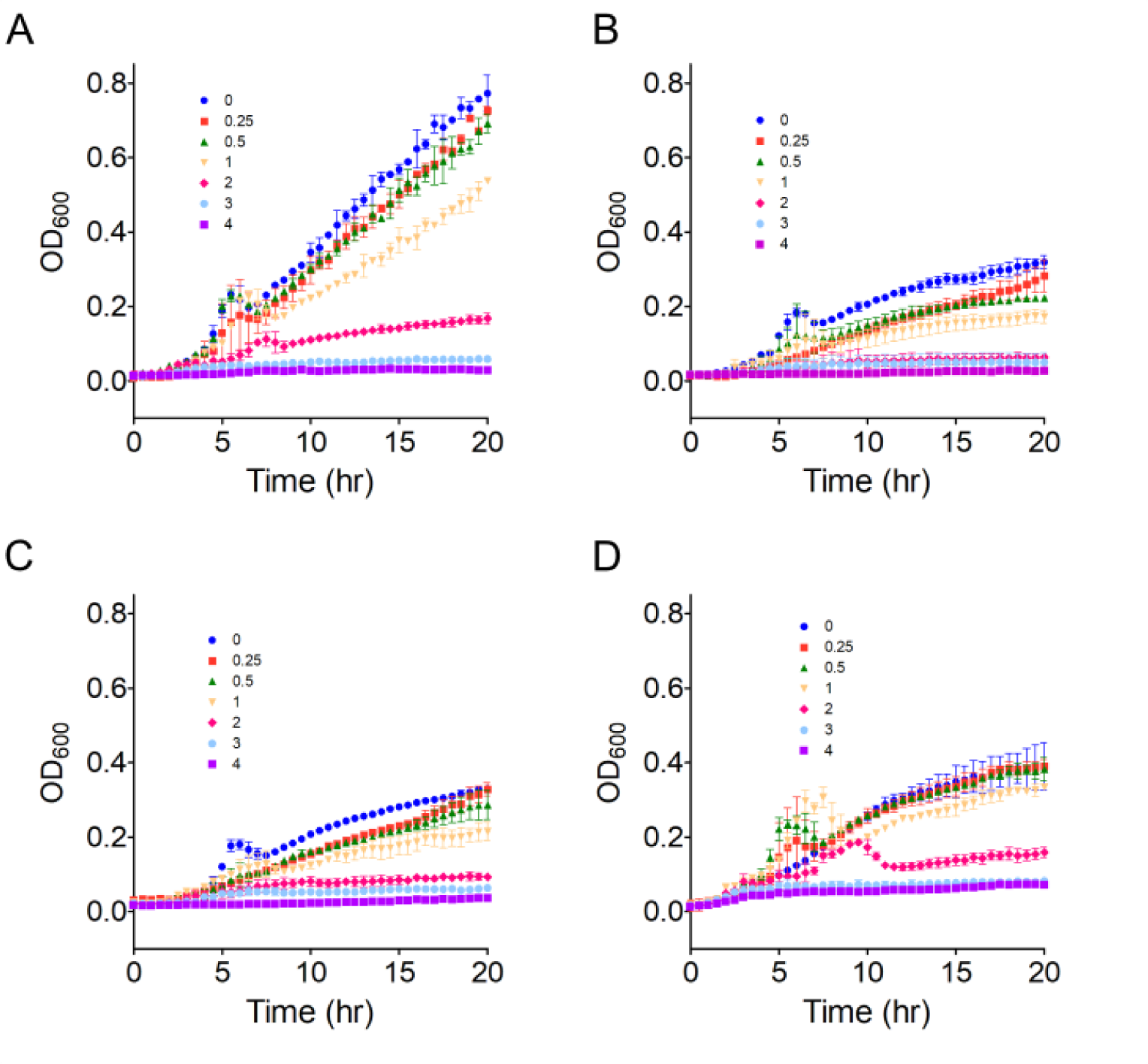
Copper toxicity in *P. aeruginosa* overexpressing OprC. Anaerobic growth of (A) pHERD30 and pHERD30-overexpressed (B) OprC_WT_, (C) OprCC143A and (D) OprC_AA_ in PA14 *ΔoprC* was monitored during copper stress in rich media supplemented with 100 mM sodium nitrate. Overexpression was induced with 0.1 % arabinose. Values indicate externally added copper in mM.

**Figure S5.**
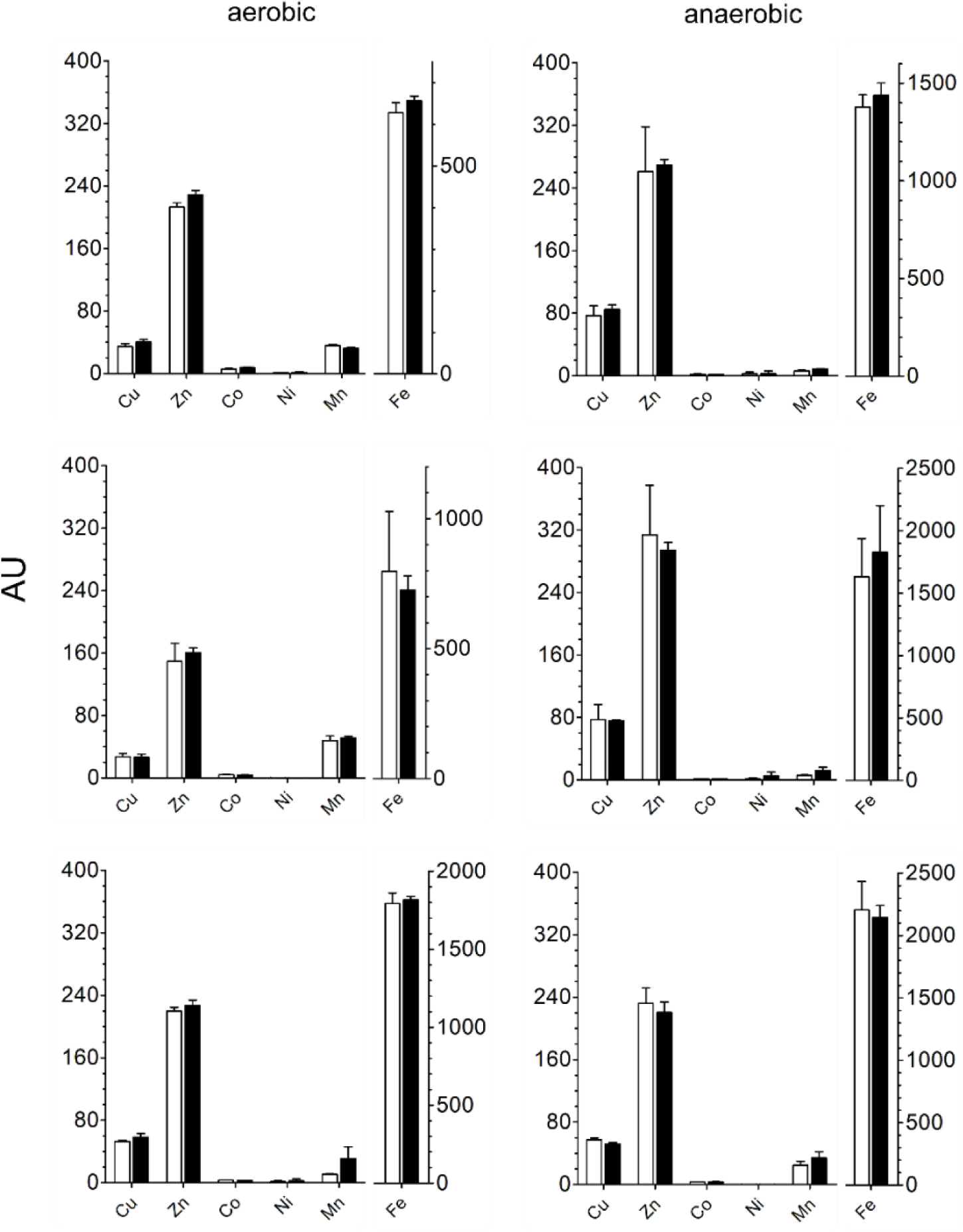
Whole cell metal content of PA14 WT and PA14 *ΔoprC* analysed via ICP-MS. Cell-associated metal content was determined in cells grown in rich media supplemented with 100 mM sodium nitrate under both aerobic (left panels) and anaerobic conditions (right panels) without added copper. The three biological replicates have been plotted separately due to the different absolute metal contents. Reported values are averages ± s.d. (n = 3).

**Figure S6.**
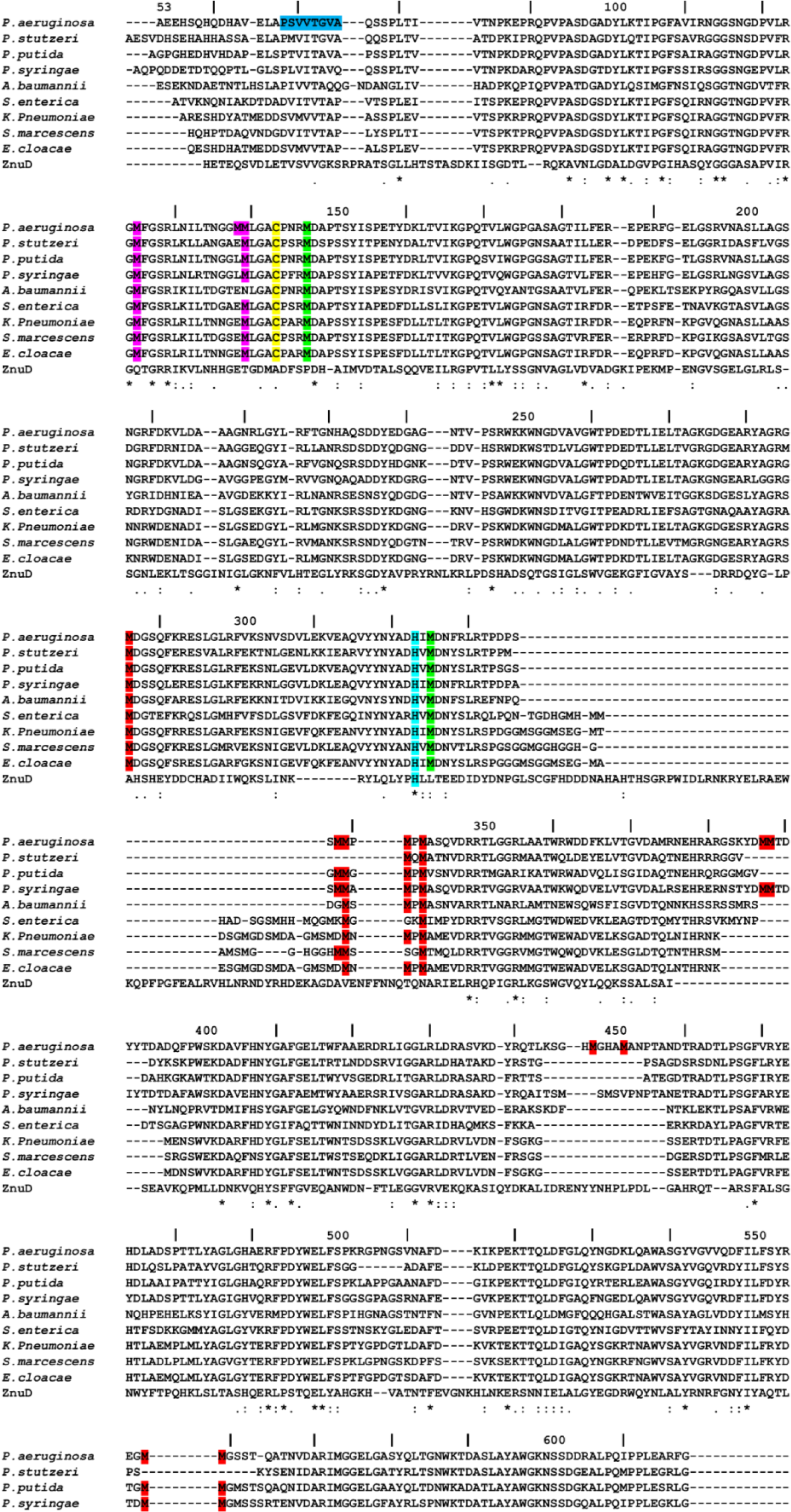
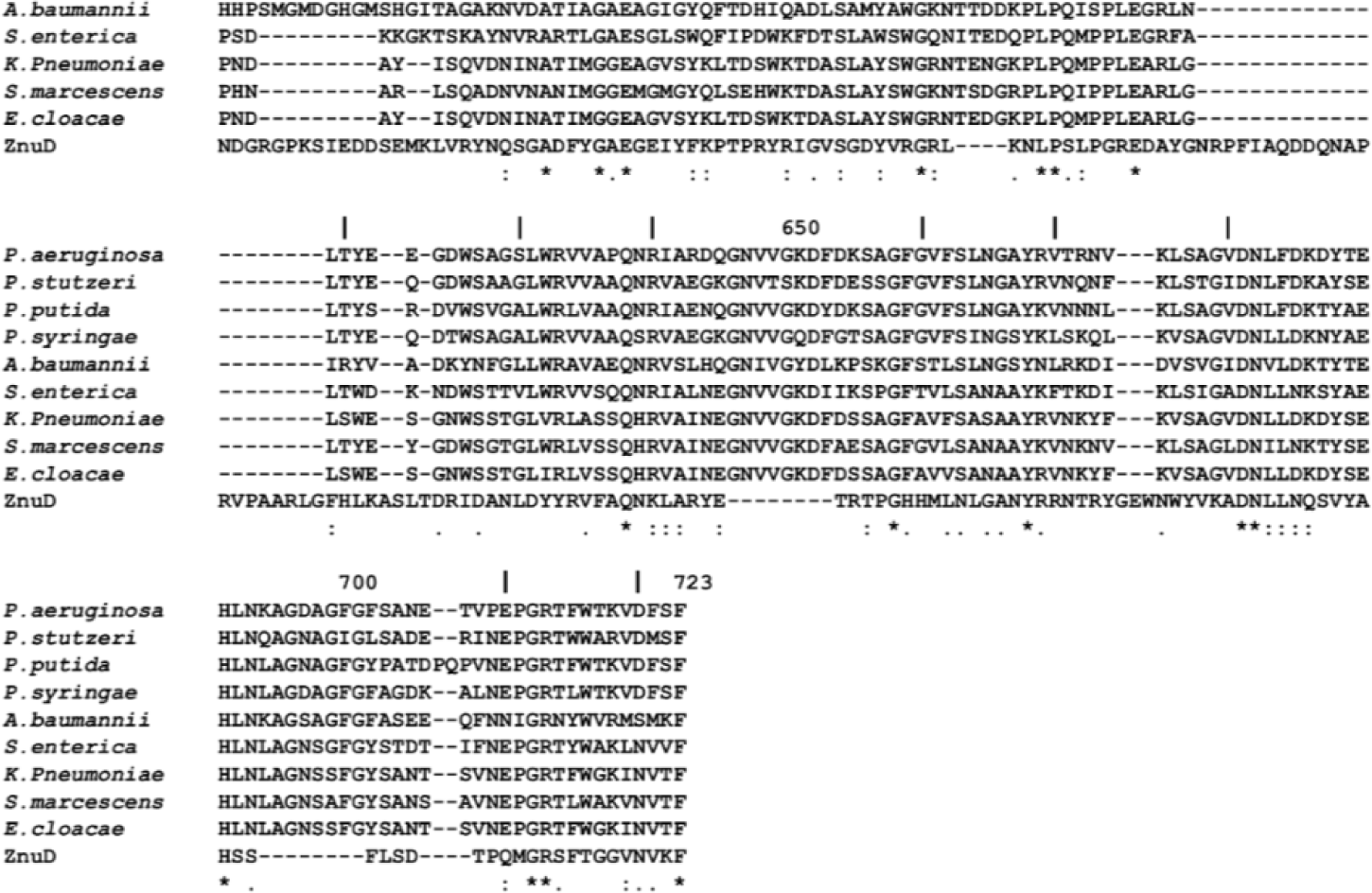
Amino acid sequence alignment for mature OprC sequences from *Pseudomonas aeruginosa* (uniprot ID G3XD89), NosA from *P. stutzeri* (uniprot ID Q00620), *P. putida* (uniprot ID Q88DI7)*, P. syringae* (uniprot ID A0A085VGG7)*, Acinetobacter baumannii* (uniprot ID A0A0G4QL30)*, Salmonella enterica* (uniprot ID A0A505CFK3)*, Klebsiella pneumonia* (uniprot ID A0A486MDQ0)*, Serratia marcescens* (uniprot ID A0A221DQ80) and *Enterobacter cloacae* (uniprot ID A0A1S6XXV6), showing high conservation of the binding site residues Cys143 (highlighted in yellow), Met147 and Met325 (green) and His323 (cyan). Methionine track residues are depicted in red, and those located in the N-terminal plug are coloured magenta. The TonB box sequence is depicted in blue. The zinc transporter ZnuD from *Neisseria meningitides* (uniprot ID Q9JZN9) is shown for comparison. Numbering is for the full-length *P. aeruginosa* OprC sequence. Clustal scoring is indicated below the alignment.

**Figure S7.**
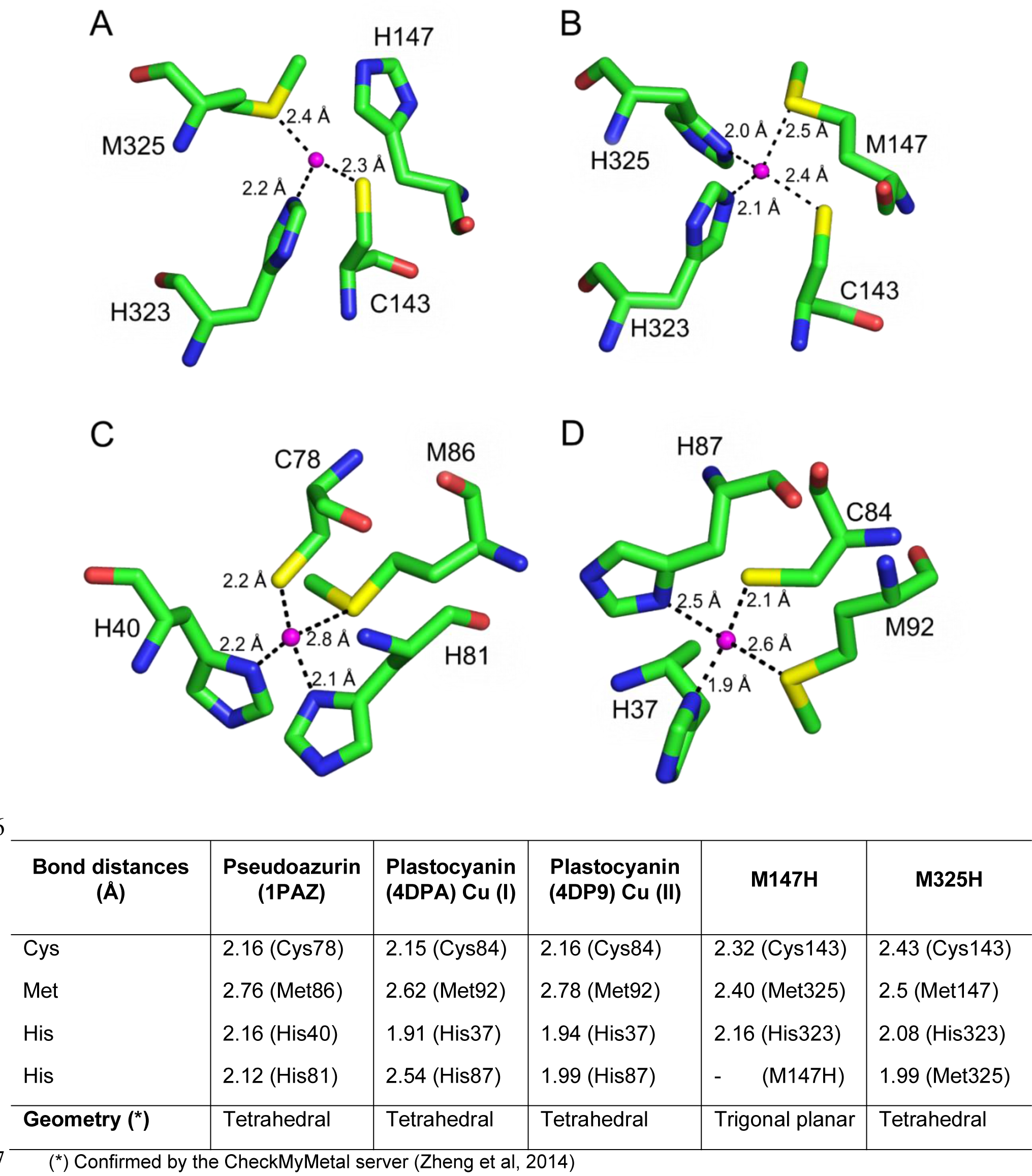
Comparison of M147H and M325H binding site residues with pseudoazurin and plastocyanin. Close up views of copper binding site residues in (A) M147H, (B) M325H, (C) pseudoazurin (PDB ID 1PAZ) and (D) plastocyanin (PDB ID 4DPA). The bound form of copper is Cu (I). Distances between coordinating residues and metal (magenta) are shown. The Table summarises distances between copper and co-ordinating residues as well as geometry.

**Figure S8.**
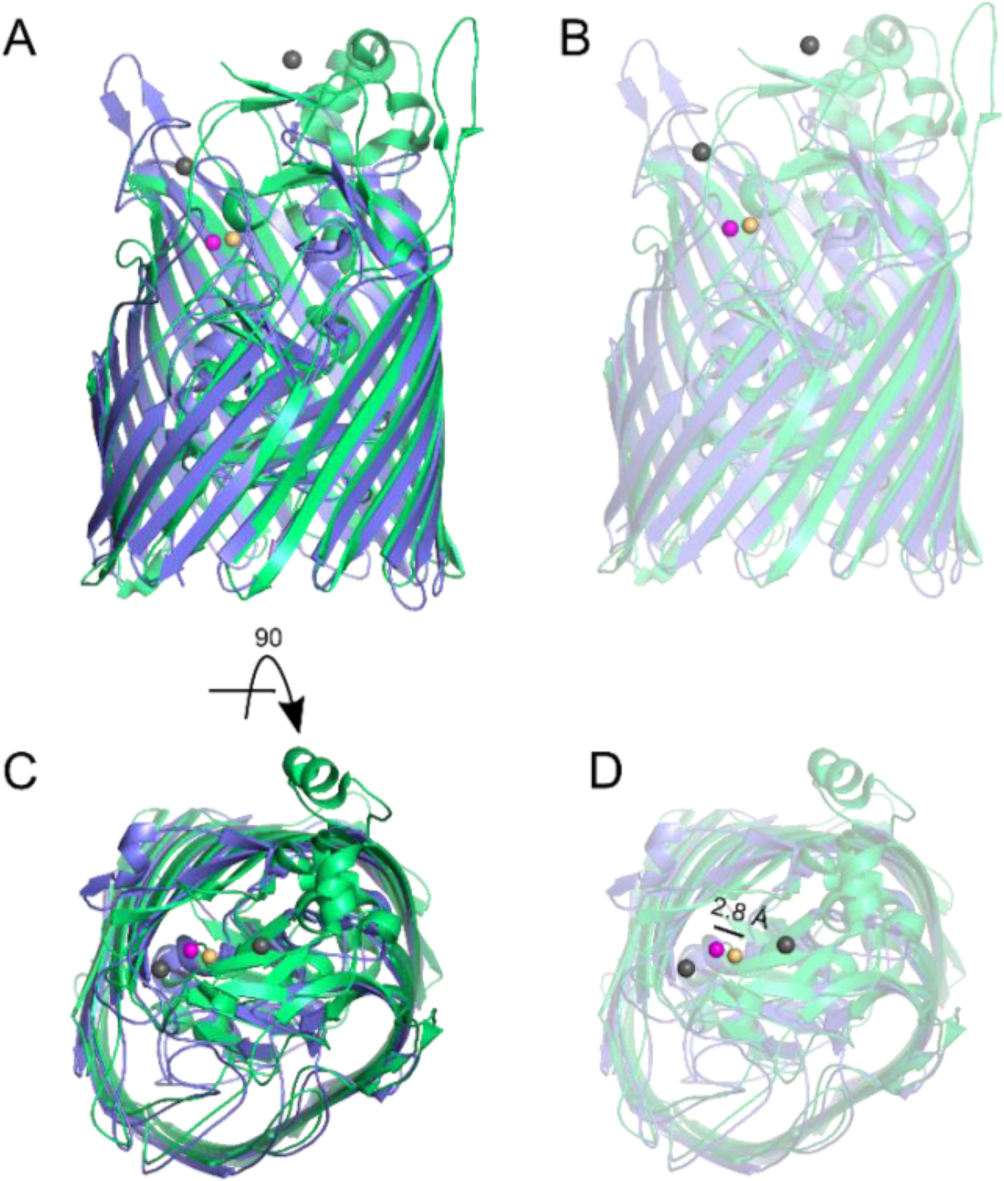
Differences between OprC and the Zn-specific ZnuD. (A, C) Cartoon representation comparing Cu-loaded OprC (coloured blue, copper atom shown as magenta sphere) and the locked version of ZnuD (coloured green, 2 cadmium atoms bound to low affinity sites represented as grey spheres, zinc bound to the high affinity site represented in orange; PDB ID 4RDR). The locked conformation of ZnuD shows low-affinity metal sites at external loops, at regions similar to the methionine track (L5) from OprC. (B, D) Transparent view of the secondary structures illustrates the similar topological location for the high affinity metal sites (distance of 2.8 Å).

**Table S1.**
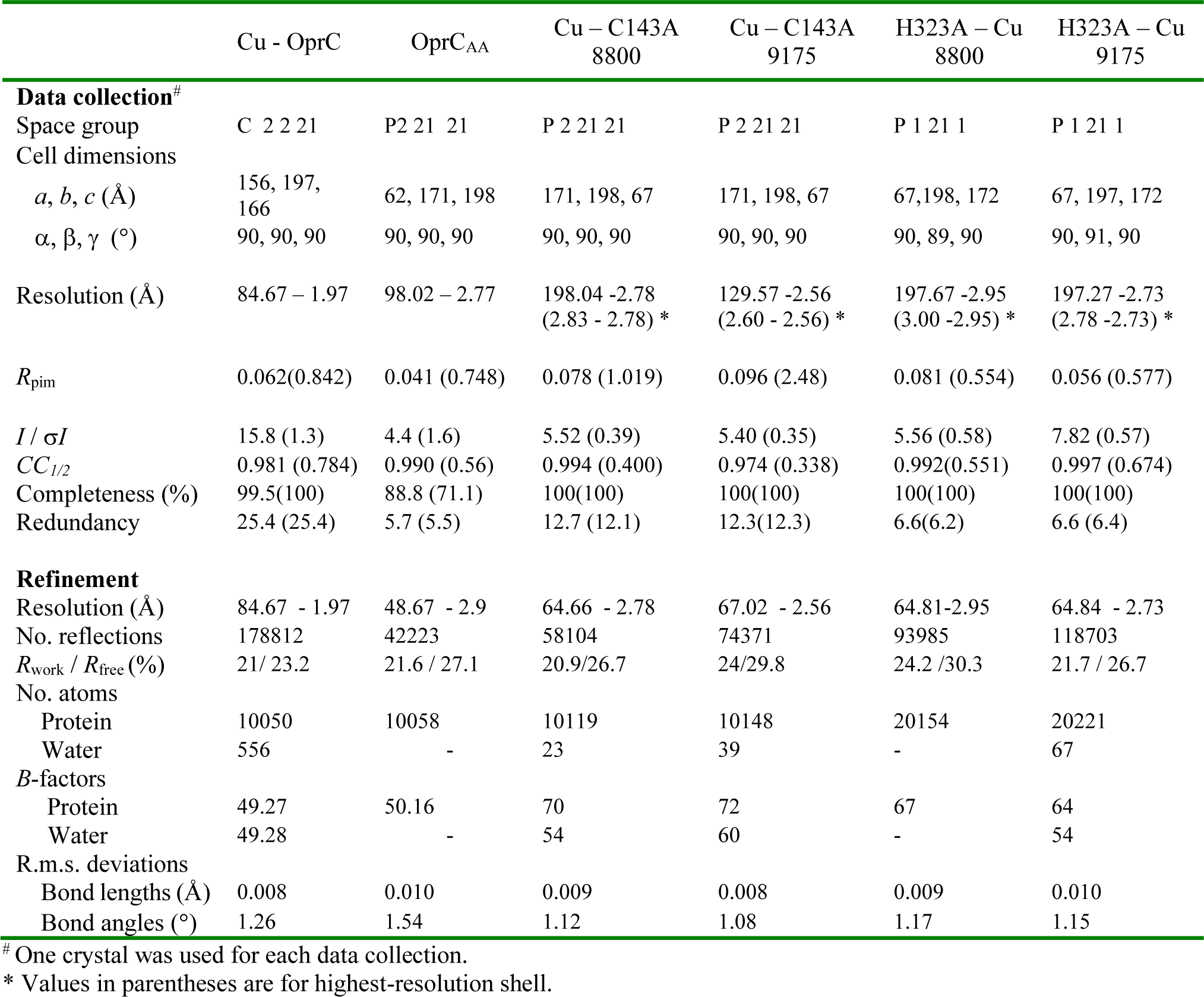
Data collection and refinement statistics for OprC variants with and without copper.

**Table S2.**
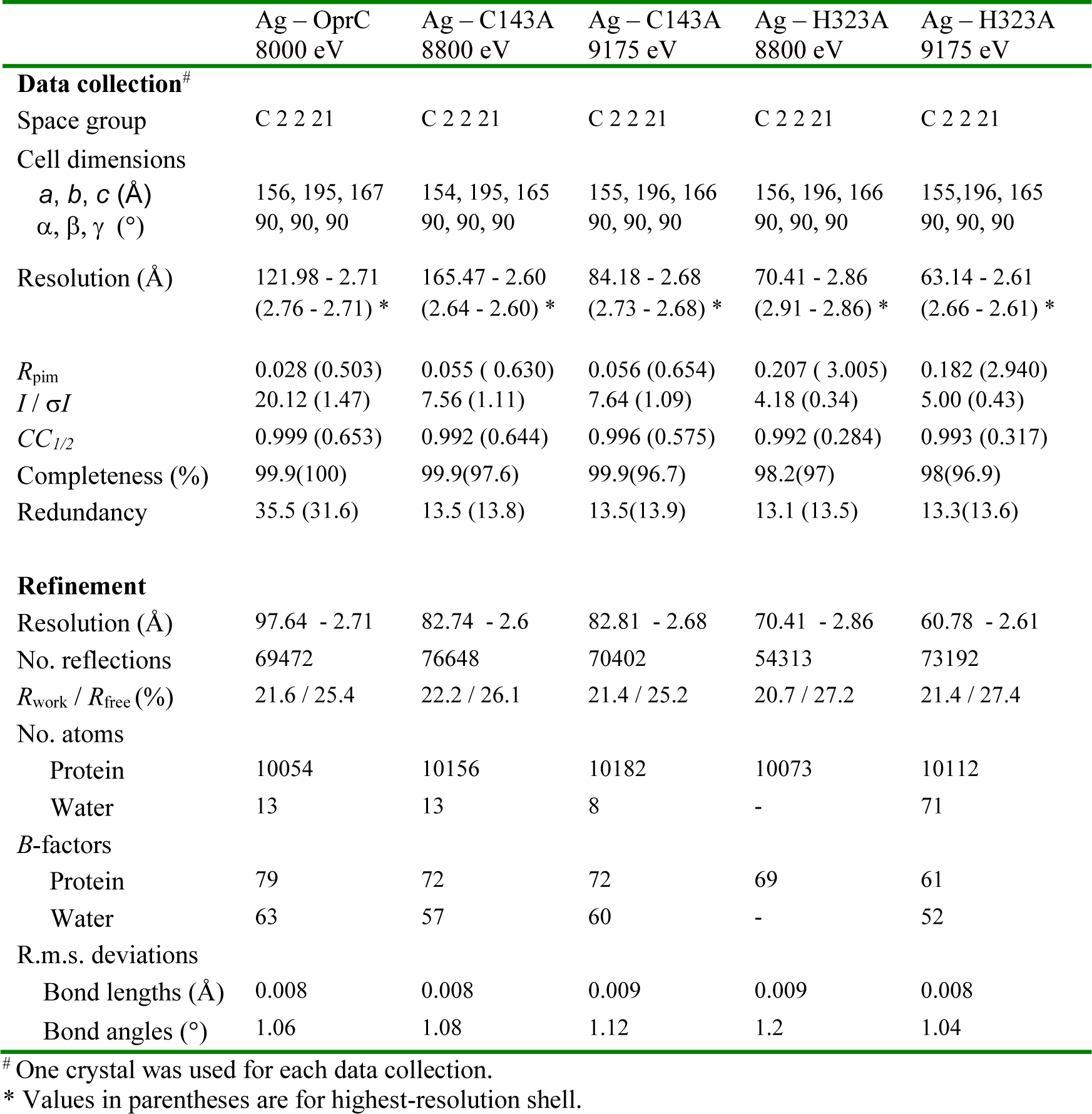
Data collection and refinement statistics for OprC variants with silver.

**Table S3.**
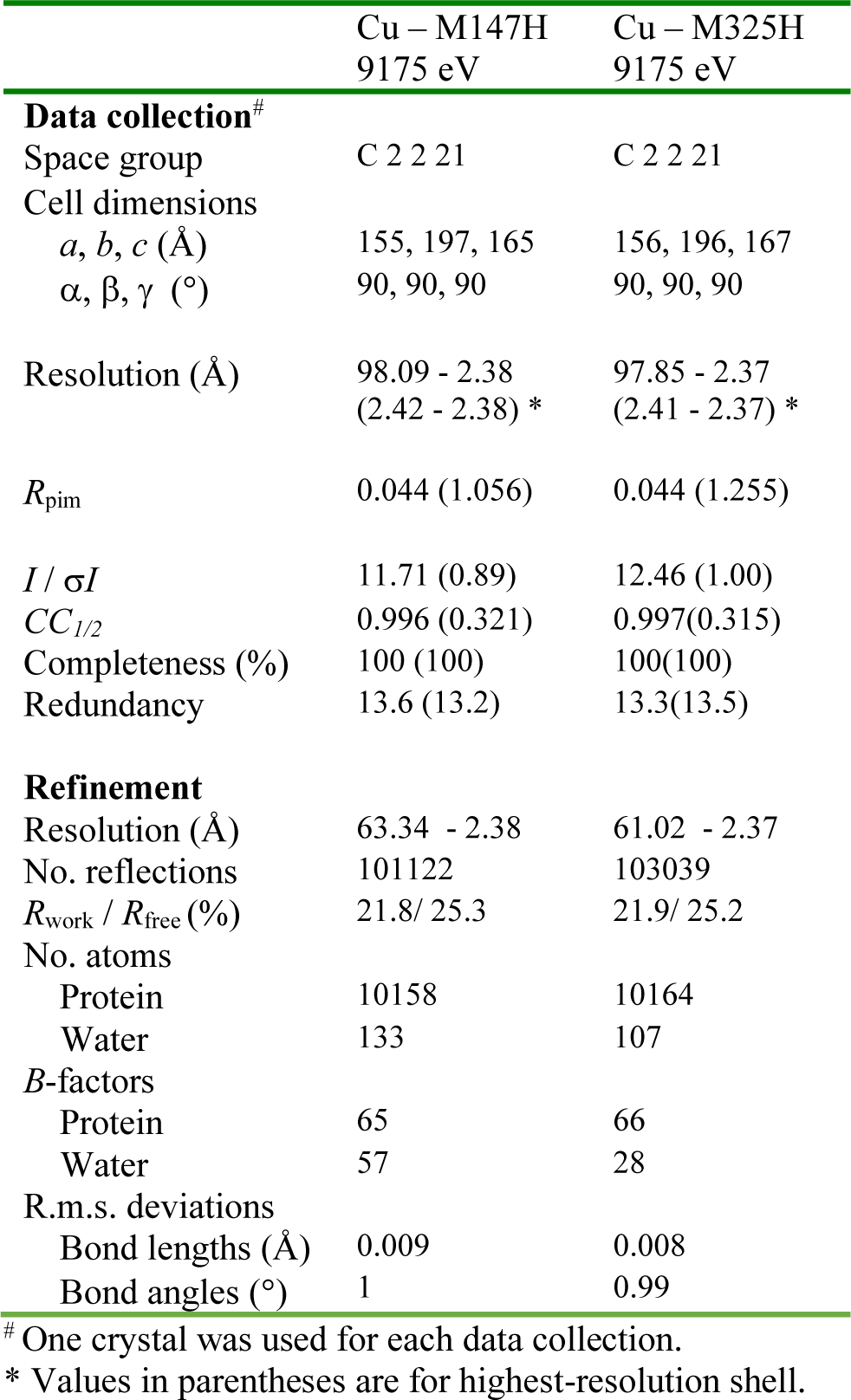
Data collection and refinement statistics for M147H and M325H variants.

## Notes

### Competing Interest Statement

The authors have declared no competing interest.

