## Supporting Information for "Acquisition of ionic copper by a bacterial outer membrane protein"

### Supplementary Information

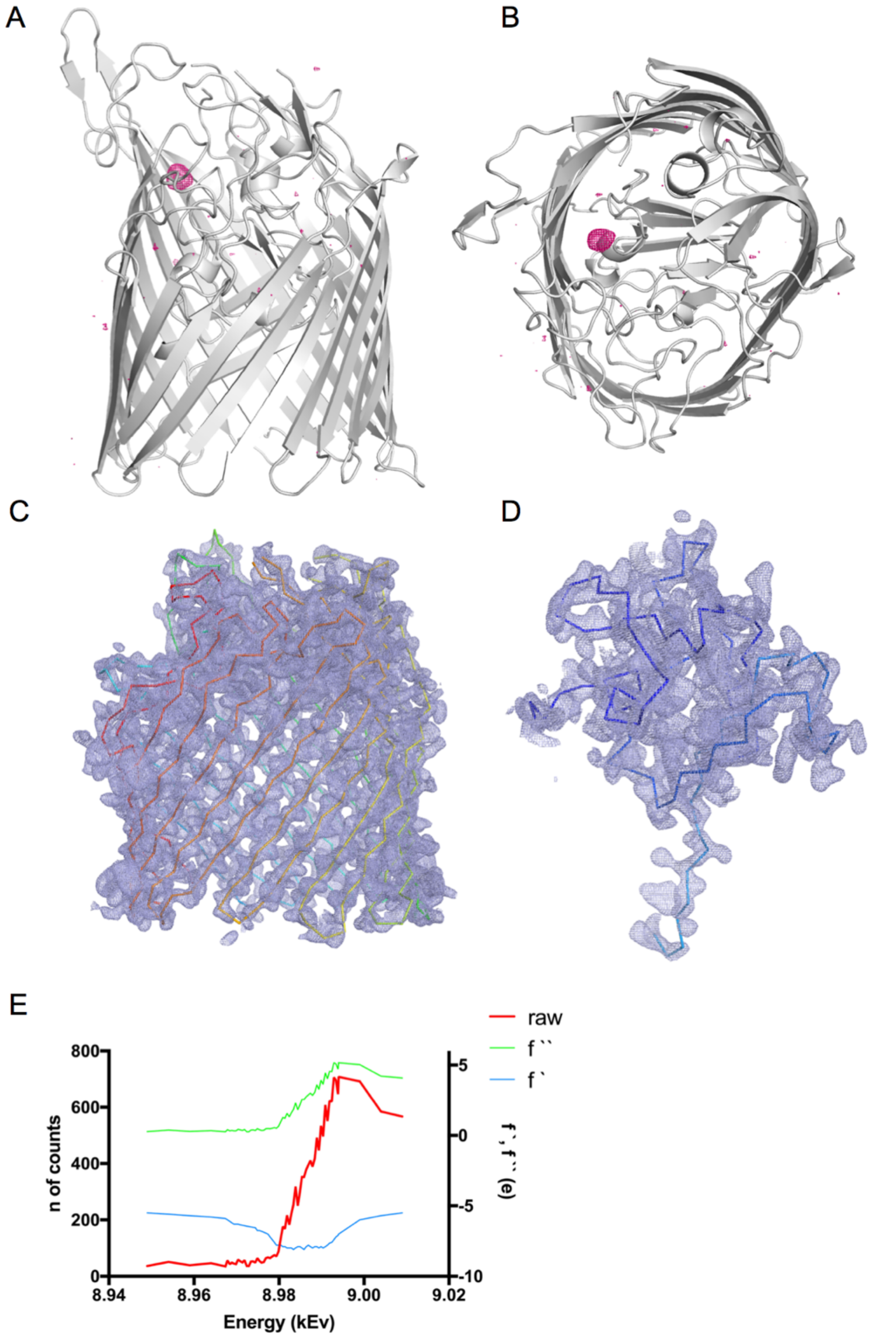


Figure S1. Anomalous data for the OprC$_{WT}$ Cu-SAD experiment. **(**A, B**)** Copper anomalous maps (coloured magenta) contoured at 4 $\sigma$ (carve = 30). Experimental density for one OprC protomer after density modification (but before model building) for (C) barrel and (D) N-terminal plug domain (map contoured at 1.5 $\sigma$, carve = 2.0). Ribbon is shown for orientation purposes. (E) X-ray fluorescence spectrum showing the copper-specific energy peak.

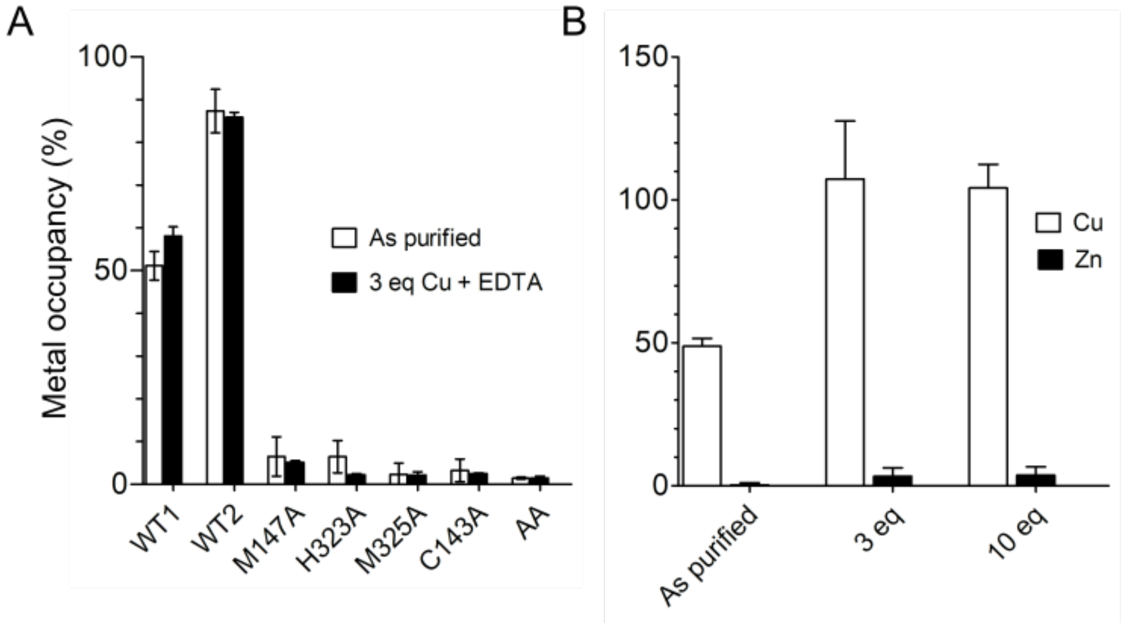


Figure S2. ICP-MS data of OprC and mutant proteins. (A) Metal occupancy of OprC and mutant proteins after incubation with 3 equivalents copper in the presence of 0.5 mM EDTA (~50-fold excess) followed by analytical size exclusion chromatography and subsequent metal analysis by ICP-MS. (B) Metal occupancy of OprC WT after incubation with 3 or 10 Eq. of Cu or Zn.

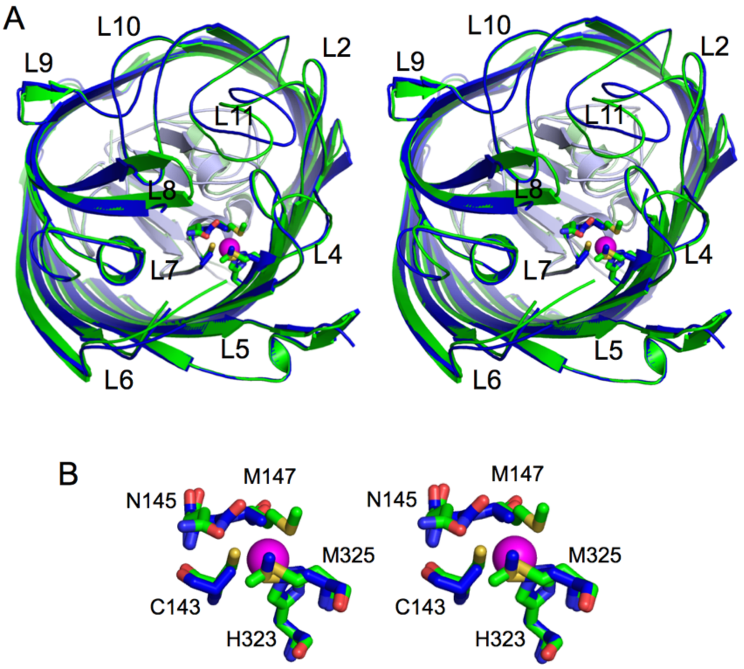


Figure S3. Stereo 3D representation of superposed OprC (coloured green) and $OprC_{AA}$ (blue). (A) Extracellular view showing loops 2, 4, 5, 6, 7, 8, 9, 10, 11 (L2, L4, L5, L6, L7, L8, L9, L10, L11). Conformational changes are observed for external loops L8 and L11. (B) Active site view illustrating superposed residues involved in metal coordination for wild type OprC (green sticks) and $OprC_{AA}$ (blue sticks). Asn145 (N145) is also shown due to its role in shielding the active site. Oxygen atoms in amino acid residues are coloured red, nitrogens blue and sulphurs yellow. Copper atom is represented as a magenta sphere.

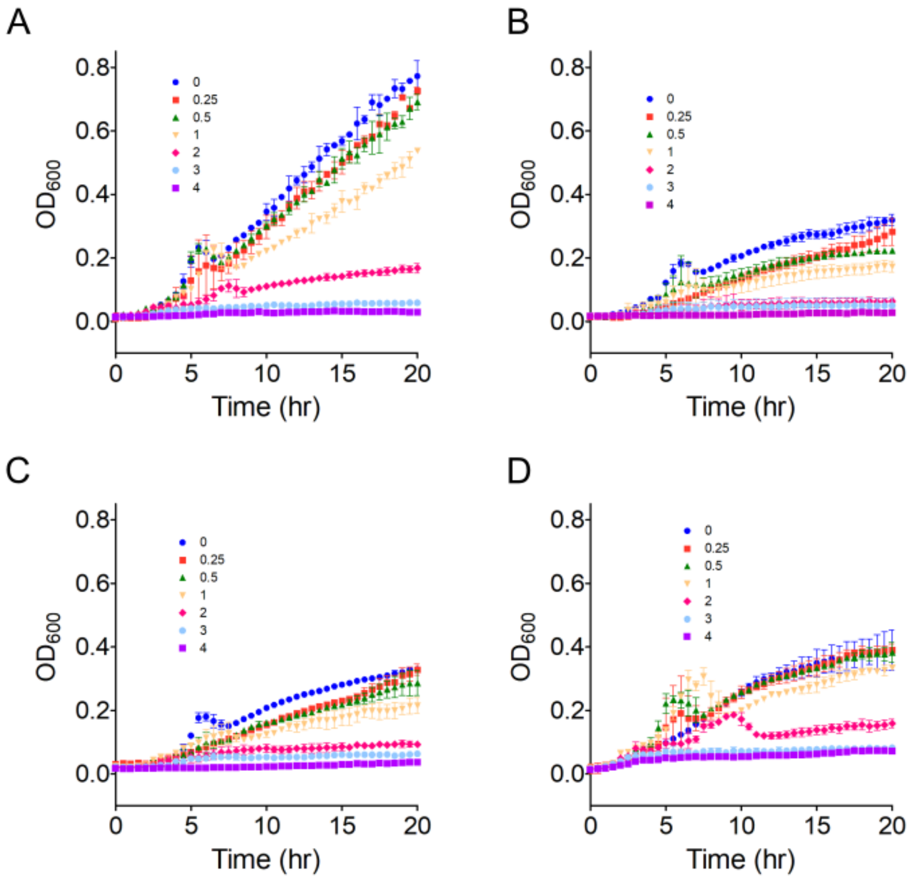


Figure S4. Copper toxicity in *P. aeruginosa* overexpressing OprC. Anaerobic growth of (A) pHERD30 and pHERD30-overexpressed (B) $\text{OprC}_{WT}$, (C) $\text{OprC}_{C143A}$ and (D) $\text{OprC}_{AA}$ in PA14 *ΔoprC* was monitored during copper stress in rich media supplemented with 100 mM sodium nitrate. Overexpression was induced with 0.1 % arabinose. Values indicate externally added copper in mM.

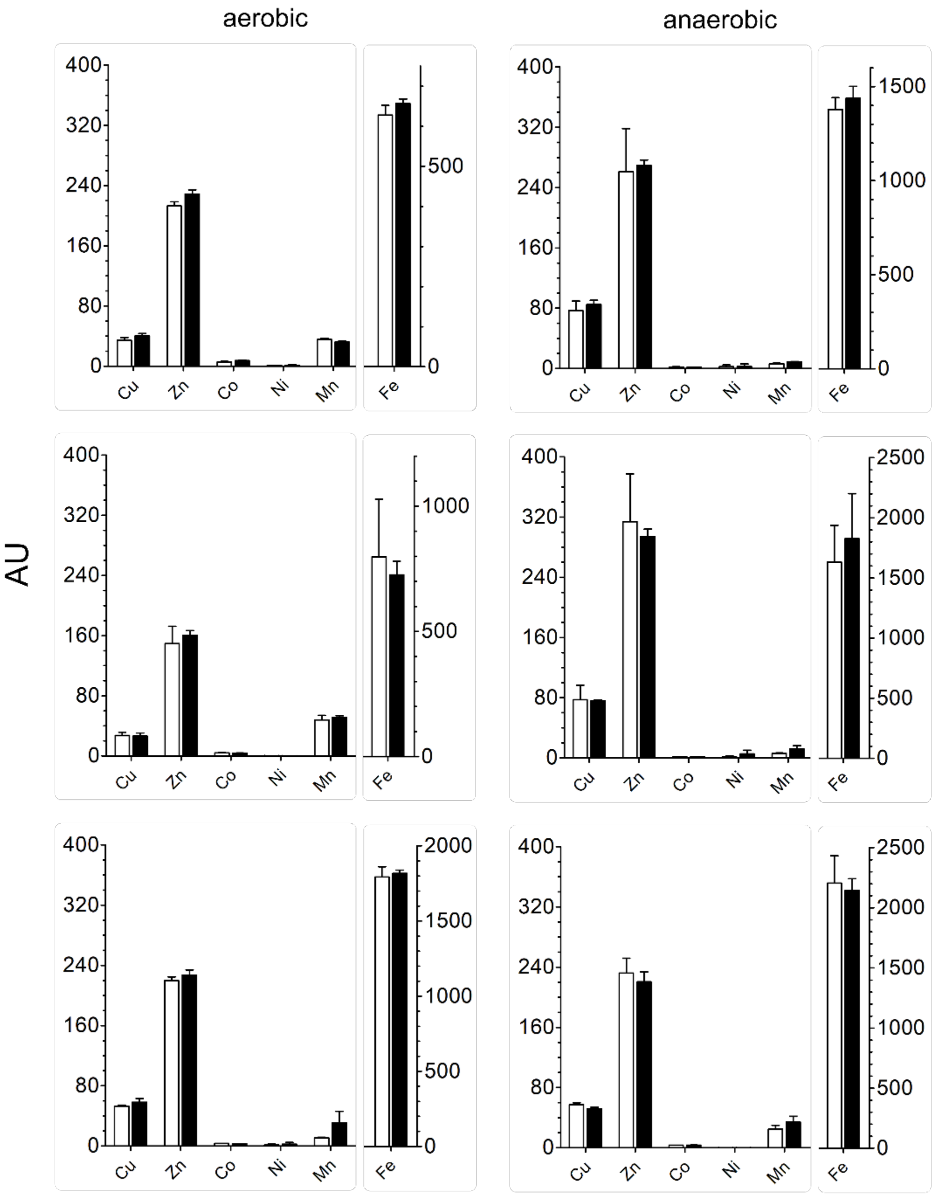


Figure S5. Whole cell metal content of PA14 WT and PA14 *ΔoprC* analysed via ICP-MS. Cell-associated metal content was determined in cells grown in rich media supplemented with 100 mM sodium nitrate under both aerobic (left panels) and anaerobic conditions (right panels) without added copper. The three biological replicates have been plotted separately due to the different absolute metal contents. Reported values are averages ± s.d. (n = 3).

```
                    53     |          |            |                |         100        |         |
P.aeruginosa    ----AEEHSQHQDHAV-ELAPSVVTGVA---QSSPLTI-------VTNPKEPRQPVPASDGADYLKTIPGFAVIRNGGSNGDPVLR
P.stutzeri      AESVDHSEHAHHASSA-ELAPMVITGVA---QQSPLTV-------ATDPKIPRQPVPASDAGDYLQTIPGFSAVRGGGSNSDPVFR
P.putida        ---AGPGHEDHVHDAP-ELSPTVITAVA---PSSPLTV-------VTNPKDPRQPVPASDGADYLKTIPGFSAIRAGGTNGDPVLR
P.syringae      -AQPQDDETDTQQPTL-GLSPLVITAVQ---QSSPLTV-------VTNPKDARQPVPASDGTDYLKTIPGFSSIRSGGSNGEPVLR
A.baumannii     ----ESEKNDAETNTLHSLAPIVVTAQQG-NDANGLIV-------HADPKQPIQPVPATDGADYLQSIMGFNSIQSGGTNGDVTFR
S.enterica      ------ATVKNQNIAKDTDADVITVTAP---VTSPLEI-------ITSPKEPRQPVPASDGSDYLKTIPGFSQIRNGGTNGDPVFR
K.Pneumoniae    -------ARESHDYATMEDDSVMVVTAP---ASSPLEV-------VTSPKRPRQPVPASDGSDYLKTIPGFSQIRAGGTNGDPVFR
S.marcescens    --------HQHPTDAQVNDGDVITVTAP---LYSPLTI-------VTSPKTPRQPVPASDGSDYLKTIPGFSQIRNGGTNGDPVFR
E.cloacae       --------QESHDHATMEDDSVMVVTAP---ALSPLEV-------VTSPKRPRQPVPASDGSDYLKTIPGFSQIRAGGTNGDPVFR
ZnuD            ----------HETEQSVDLETVSVVGKSRPRATSGLLHTSTASDKIISGDTL--RQKAVNLGDALDGVPGIHASQYGGGASAPVIR
                                        .        . *           . .       * :  * *. : *:   : **  .  .:*

                      |         |        150        |          |          |            |        200
P.aeruginosa    GMFGSRLNILTNGGMMLGACPNRMDAPTSYISPETYDKLTVIKGPQTVLWGPGASAGTILFER--EPERFG-ELGSRVNASLLAGS
P.stutzeri      GMFGSRLKLLANGAEMLGACPSRMDSPSSYITPENYDALTVIKGPQTVLWGPGNSAATILLER--DPEDFS-ELGGRIDASFLVGS
P.putida        GMFGSRLNILTNGGLMLGACPNRMDAPTSYISPETYDRLTVIKGPQSVIWGPGGSAGTILFER--EPEKFG-TLGSRVNASLLAGS
P.syringae      GMFGSRLNLRTNGGLMLGACPFRMDAPSSYIAPETFDKLTVVKGPQTVQWGPGASAGTVLFER--EPEHFG-ELGSRLNGSVLAGS
A.baumannii     GMFGSRIKILTDGTENLGACPNRMDAPTSYISPESYDRISVIKGPQTVQYANTGSAATVLFER--QPEKLTSEKPYRGQASVLLGS
S.enterica      GMFGSRLKILTDGAEMLGACPSRMDAPTSYIAPEDFDLLSLIKGPETVLWGPGNSAGTIRFDR--ETPSFE-TNAVKGTASVLAGS
K.Pneumoniae    GMFGSRLRILTNNGEMLGACPARMDAPSSYISPESFDLLTLTKGPQTVLWGPGNSAGTIRFDR--EQPRFN-KPGVQGNASLLAAS
S.marcescens    GMFGSRLKILTDGSEMLGACPSRMDAPTSYISPESFDLLTITKGPQTVLWGPGSSAGTVRFER--ERPRFD-KPGIKGSASVLTGS
E.cloacae       GMFGSRLRILTNNGEMLGACPARMDAPSSYISPESFDLLTLTKGPQTVLWGPGNSAGTIRFDR--EQPRFD-KPGVQGNASLLAAS
ZnuD            GQTGRRIKVLNHHGETGDMADFSPDH-AIMVDTALSQQVEILRGPVTLLYSSGNVAGLVDVADGKIPEKMP-ENGVSGELGLRLS-
                *  * *:.:  .     . .    *  :  :         : : : :** :: :.    *. : .        :          ..

                   |           |          |         |          250            |         |         |
P.aeruginosa    NGRFDKVLDA--AAGNRLGYL-RFTGNHAQSDDYEDGAG---NTV-PSRWKKWNGDVAVGWTPDEDTLIELTAGKGDGEARYAGRG
P.stutzeri      DGRFDRNIDA--AAGGEQGYI-RLLANRSDSDDYQDGNG---DDV-HSRWDKWSTDLVLGWTPDEDTLLELTVGRGDGEARYAGRM
P.putida        NGRFDKVLDA--AAGNSQGYA-RFVGNQSRSDDYHDGNK---DTV-PSRWEKWNGDVALGWTPDQDTLLELTAGKGDGEARYAGRG
P.syringae      NGRFDKVLDG--AVGGPEGYM-RVVGNQAQADDYKDGRG---NTV-PSRWEKWNGDVALGWTPDADTLIELTAGKGNGEARLGGRG
A.baumannii     YGRIDHNIEA--AVGDEKKYI-RLNANRSESNSYQDGDG---NTV-PSAWKKWNVDVALGFTPDENTWVEITGGKSDGESLYAGRS
S.enterica      RDRYDGNADI--SLGSEKGYL-RLTGNKSRSSDYKDGNG---KNV-HSGWDKWNSDITVGITPEADRLIEFSAGTGNAQAAYAGRA
K.Pneumoniae    NNRWDENADI--SLGSEDGYL-RLMGNKSRSDDYKDGNG---DRV-PSKWDKWNGDMALGWTPDKDTLIELTAGKGDGESRYAGRS
S.marcescens    NGRWDENIDA--SLGAEQGYL-RVMANKSRSNDYQDGTN---TRV-PSRWDKWNGDLALGWTPDNDTLLEVTMGRGNGEARYAGRS
E.cloacae       KNRWDENADI--SLGSEDGYL-RLMGNKSRSDDYKDGNG---DRV-PSKWDKWNGDMALGWTPDKDTLIELTAGKGDGESRYAGRS
ZnuD            SGNLEKLTSGGINIGLGKNFVLHTEGLYRKSGDYAVPRYRNLKRLPDSHADSQTGSIGLSWVGEKGFIGVAYS---DRRDQYG-LP
                 .. :   .     *    :  :  .    :..*          :  *  .. . .: :. . : .          : .   .

                    |        300          |          |         |
P.aeruginosa    MDGSQFKRESLGLRFVKSNVSDVLEKVEAQVYYNYADHIMDNFRLRTPDPS-----------------------------------
P.stutzeri      MDGSQFERESVALRFEKTNLGENLKKIEARVYYNYADHVMDNYSLRTPPM------------------------------------
P.putida        MDGSQFKRESLGLRFEKSNLGEVLDKVEAQVYYNYADHVMDNYSLRTPSGS-----------------------------------
P.syringae      MDSSQLERESLGLKFEKRNLGGVLDKLEAQVYYNYADHIMDNFRLRTPDPA-----------------------------------
A.baumannii     MDGSQFARESLGLRFEKKNITDVIKKIEGQVNYSYNDHVMDNFSLREFNPQ-----------------------------------
S.enterica      MDGTEFKRQSLGMHFVFSDLGSVFDKFEGQINYNYARHVMDNYSLRQLPQN-TGDHGMH-MM------------------------
K.Pneumoniae    MDGSQFRRESLGARFEKSNIGEVFQKFEANVYYNYADHIMDNYSLRSPDGGMSGGMSEG-MT------------------------
S.marcescens    MDGSQFKRESLGMRVEKSNIGEVLDKLEAQVYYNYANHVMDNVTLRSPGSGGMGGHGGH-G-------------------------
E.cloacae       MDGSQFSRESLGARFGKSNIGEVFQKFEANVYYNYADHIMDNYSLRSPGGGMSGGMSEG-MA------------------------
ZnuD            AHSHEYDDCHADIIWQKSLINK--------RYLQLYPHLLTEEDIDYDNPGLSCGFHDDDNAHAHTHSGRPWIDLRNKRYELRAEW
                 .. :              :             .   *:: :                      :

                                           |              350         |         |         |     |
P.aeruginosa    --------------------------SMMP------MPMASQVDRRTLGGRLAATWRWDDFKLVTGVDAMRNEHRARGSKYDMMTD
P.stutzeri      ------------------------------------MQMATNVDRRTLGGRMAATWQLDEYELVTGVDAQTNEHRRRGGV------
P.putida        --------------------------GMMG------MPMVSNVDRRTMGARIKATWRWADVQLISGIDAQTNEHRQRGGMGV----
P.syringae      --------------------------SMMA------MPMASQVDRRTVGGRVAATWKWQDVELVTGVDALRSEHRERNSTYDMMTD
A.baumannii     --------------------------DGMS------MPMASNVARRTLNARLAMTNEWSQWSFISGVDTQNNKHSSRSSMRS----
S.enterica      ------------HAD-SGSMHH-MQGMKMG------GKMIMPYDRRTVSGRLMGTWDWEDVKLEAGTDTQMYTHRSVKMYNP----
K.Pneumoniae    ------------DSGMGDSMDA-GMSMDMN------MPMAMEVDRRTVGGRMMGTWEWADVELKSGADTQLNIHRNK---------
S.marcescens    ------------AMSMG----G-HGGHMMS------SGMTMQLDRRTVGGRVMGTWQWQDVKLESGLDTQTNTHRSM---------
E.cloacae       ------------ESGMGDSMDA-GMSMDMN------MPMAMEVDRRTVGGRMMGTWEWADVELKSGADTQLNTHRNK---------
ZnuD            KQPFPGFEALRVHLNRNDYRHDEKAGDAVENFFNNQTQNARIELRHQPIGRLKGSWGVQYLQQKSSALSAI---------------
                                                            *:   .*:  :      .  :.  :

                          400       |         |         |          |          450        |         |
P.aeruginosa    YYTDADQFPWSKDAVFHNYGAFGELTWFAAERDRLIGGLRLDRASVKD-YRQTLKSG--HMGHAMANPTANDTRADTLPSGFVRYE
P.stutzeri      ---DYKSKPWEKDADFHNYGLFGELTRTLNDDSRVIGGARLDHATAKD-YRSTG-------------PSAGDSRSDNLPSGFLRYE
P.putida        --DAHKGKAWTKDADFHNYGAFSELTWYVSGEDRLITGARLDRASARD-FRTTS-------------ATEGDTRADTLPSGFIRYE
P.syringae      IYTDTDAFAWSKDAVEHNYGAFAEMTWYAAERSRIVSGARLDRASAKD-YRQAITSM----SMSVPNPTANETRADTLPSGFARYE
A.baumannii     ---NYLNQPRVTDMIFHSYGAFGELGYQWNDFNKLVTGVRLDRVTVED-ERAKSKDF-------------NTKLEKTLPSAFVRWE
S.enterica      --DTSGAGPWNKDARFHDYGIFAQTTWNINNDYDLITGARIDHAQMKS-FKKA-----------------ERKRDAYLPAGFVRTE
K.Pneumoniae    -----MENSWVKDARFHDYGLFSELTWNTSDSSKLVGGARLDRVLVDN-FSGKG----------------SSERTDTLPAGFVRFE
S.marcescens    -----SRGSWEKDAQFNSYGAFSELTWSTSEQDKLIGGARLDRTLVEN-FRSGS----------------DGERSDTLPSGFMRLE
E.cloacae       -----MDNSWVKDARFHDYGLFSELTWNTSDSSKLVGGARLDRVLVDN-FSGKG----------------SSERTDTLPAGFVRFE
ZnuD            --SEAVKQPMLLDNKVQHYSFFGVEQANWDN-FTLEGGVRVEKQKASIQYDKALIDRENYYNHPLPDL--GAHRQT--ARSFALSG
                            *   : *. *.           :  * *:::                                      .*

                      |         |        500        |             |         |         |        550
P.aeruginosa    HDLADSPTTLYAGLGHAERFPDYWELFSPKRGPNGSVNAFD----KIKPEKTTQLDFGLQYNGDKLQAWASGYVGVVQDFILFSYR
P.stutzeri      HDLQSLPATAYVGLGHTQRFPDYWELFSGG------ADAFE----KLDPEKTTQLDFGLQYSKGPLDAWVSAYVGQVRDYILFSYS
P.putida        HDLAAIPATTYIGLGHAQRFPDYWELFSPKLAPPGAANAFD----GIKPEKTTQLDFGIQYRTERLEAWASGYVGQIRDYILFDYR
P.syringae      YDLADSPTTLYAGIGHVQRFPDYWELFSGGSGPAGSRNAFE----GVKPEKTTQLDFGAQFNGEDLQAWVSGYVGQVRDFILFDYS
A.baumannii     NQHPEHELKSYIGLGYVERMPDYWELFSPIHGNAGSTNTFN----GVNPEKTLQLDMGFQQQHGALSTWASAYAGLVDDYILMSYH
S.enterica      HTFSDKKGMMYAGLGYVKRFPDYWELFSSTNSKYGLEDAFT----SVRPEETTQLDIGTQYNIGDVTTWVSFYTAYINNYIIFQYD
K.Pneumoniae    HTLAEMPLMLYAGLGYTERFPDYWELFSPTYGPDGTLDAFD----KVKTEKTTQLDIGAQYSGKRTNAWVSAYVGRVNDFILFRYD
S.marcescens    HTLADLPLMLYAGVGYTERFPDYWELFSPKLGPNGSKDPFS----SVKSEKTTQLDIGAQYNGKRFNGWVSAYVGRVDDFILFKYD
E.cloacae       HTLAEMQLMLYAGLGYTERFPDYWELFSPTFGPDGTSDAFD----KVKTEKTTQLDIGAQYSGKRTNAWVSAYVGRVNDFILFRYD
ZnuD            NWYFTPQHKLSLTASHQERLPSTQELYAHGKH--VATNTFEVGNKHLNKERSNNIELALGYEGDRWQYNLALYRNRFGNYIYAQTL
                              .: :*:*.  **::          : *      :  *.: ::::.            : *   . ::*

                             |          |         |         |       600         |
P.aeruginosa    EGM---------MGSST-QATNVDARIMGGELGASYQLTGNWKTDASLAYAWGKNSSDDRALPQIPPLEARFG-------------
P.stutzeri      PS---------------KYSENIDARIMGGELGATYRLTSNWKTDASLAYAWGKNSSDGEALPQMPPLEGRLG-------------
P.putida        TGM---------MGMSTSQAQNIDARIMGGELGAAYQLTDNWKADATLAYAWGKNSSDGKALPQMPPLESRLG-------------
P.syringae      TDM---------MGMSSSRTENVDARIMGGELGFAYRLSPNWKTDATLAYAWGKNSSDGQALPQIPPLEGKLG-------------
```

45

```
A.baumannii     HHPSMGMDGHGMSHGITAGAKNVDATIAGAEAGIGYQFTDHIQADLSAMYAWGKNTTDDKPLPQISPLEGRLN-------------
S.enterica      PSD---------KKGKTSKAYNVRARTLGAESGLSWQFIPDWKFDTSLAWSWGQNITEDQPLPQMPPLEGRFA-------------
K.Pneumoniae    PND---------AY--ISQVDNINATIMGGEAGVSYKLTDSWKTDASLAYSWGRNTENGKPLPQMPPLEARLG-------------
S.marcescens    PHN---------AR--LSQADNVNANIMGGEMGMGYQLSEHWKTDASLAYSWGKNTSDGRPLPQIPPLEARLG-------------
E.cloacae       PND---------AY--ISQVDNINATIMGGEAGVSYKLTDSWKTDASLAYSWGRNTEDGKPLPQMPPLEARLG-------------
ZnuD            NDGRGPKSIEDDSEMKLVRYNQSGADFYGAEGEIYFKPTPRYRIGVSGDYVRGRL----KNLPSLPGREDAYGNRPFIAQDDQNAP
                                     :  *   *.*    ::     : . :  :  *:     . **.:   *

                         |            |         |         650       |         |            |
P.aeruginosa    --------LTYE--E-GDWSAGSLWRVVAPQNRIARDQGNVVGKDFDKSAGFGVFSLNGAYRVTRNV---KLSAGVDNLFDKDYTE
P.stutzeri      --------LTYE--Q-GDWSAAGLWRVVAAQNRVAEGKGNVTSKDFDESSGFGVFSLNGAYRVNQNF---KLSTGIDNLFDKAYSE
P.putida        --------LTYS--R-DVWSVGALWRLVAAQNRIAENQGNVVGKDYDKSAGFGVFSLNGAYKVNNNL---KLSAGVDNLFDKTYAE
P.syringae      --------LTYE--Q-DTWSAGALWRVVAAQSRVAEGKGNVVGQDFGTSAGFGVFSINGSYKLSKQL---KVSAGVDNLLDKNYAE
A.baumannii     --------IRYV--A-DKYNFGLLWRAVAEQNRVSLHQGNIVGYDLKPSKGFSTLSLNGSYNLRKDI---DVSVGIDNVLDKTYTE
S.enterica      --------LTWD--K-NDWSTTVLWRVVSQQNRIALNEGNVVGKDIIKSPGFTVLSANAAYKFTKDI---KLSIGADNLLNKSYAE
K.Pneumoniae    --------LSWE--S-GNWSSTGLVRLASSQHRVAINEGNVVGKDFDSSAGFAVFSASAAYRVNKYF---KVSAGVDNLLDKDYSE
S.marcescens    --------LTYE--Y-GDWSGTGLWRLVSSQHRVAINEGNVVGKDFAESAGFGVLSANAAYKVNKNV---KLSAGLDNILNKTYSE
E.cloacae       --------LSWE--S-GNWSSTGLIRLVSSQHRVAINEGNVVGKDFDSSAGFAVVSANAAYRVNKYF---KVSAGVDNLLDKDYSE
ZnuD            RVPAARLGFHLKASLTDRIDANLDYYRVFAQNKLARYE--------TRTPGHHMLNLGANYRRNTRYGEWNWYVKADNLLNQSVYA
                        :       .  .       .  * :::  :          : *.  .. .. *.        .     **::::

                         700         |         |  723
P.aeruginosa    HLNKAGDAGFGFSANE--TVPEPGRTFWTKVDFSF
P.stutzeri      HLNQAGNAGIGLSADE--RINEPGRTWWARVDMSF
P.putida        HLNLAGNAGFGYPATDPQPVNEPGRTFWTKVDFSF
P.syringae      HLNLAGDAGFGFAGDK--ALNEPGRTLWTKVDFSF
A.baumannii     HLNKAGSAGFGFASEE--QFNNIGRNYWVRMSMKF
S.enterica      HLNLAGNSGFGYSTDT--IFNEPGRTYWAKLNVVF
K.Pneumoniae    HLNLAGNSSFGYSANT--SVNEPGRTFWGKINVTF
S.marcescens    HLNLAGNSAFGYSANS--AVNEPGRTLWAKVNVTF
E.cloacae       HLNLAGNSSFGYSANT--SVNEPGRTFWGKINVTF
ZnuD            HSS--------FLSD----TPQMGRSFTGGVNVKF
                * .                  : **.    :.. *
```

Figure S6. Amino acid sequence alignment for mature OprC sequences from *Pseudomonas aeruginosa* (uniprot ID G3XD89), NosA from *P. stutzeri* (uniprot ID Q00620), *P. putida* (uniprot ID Q88DI7), *P. syringae* (uniprot ID A0A085VGG7), *Acinetobacter baumannii* (uniprot ID A0A0G4QL30), *Salmonella enterica* (uniprot ID A0A505CFK3), *Klebsiella pneumonia* (uniprot ID A0A486MDQ0), *Serratia marcescens* (uniprot ID A0A221DQ80) and *Enterobacter cloacae* (uniprot ID A0A1S6XXV6), showing high conservation of the binding site residues Cys143 (highlighted in yellow), Met147 and Met325 (green) and His323 (cyan). Methionine track residues are depicted in red, and those located in the N-terminal plug are coloured magenta. The TonB box sequence is depicted in blue. The zinc transporter ZnuD from *Neisseria meningitides* (uniprot ID Q9JZN9) is shown for comparison. Numbering is for the full-length *P. aeruginosa* OprC sequence. Clustal scoring is indicated below the alignment.

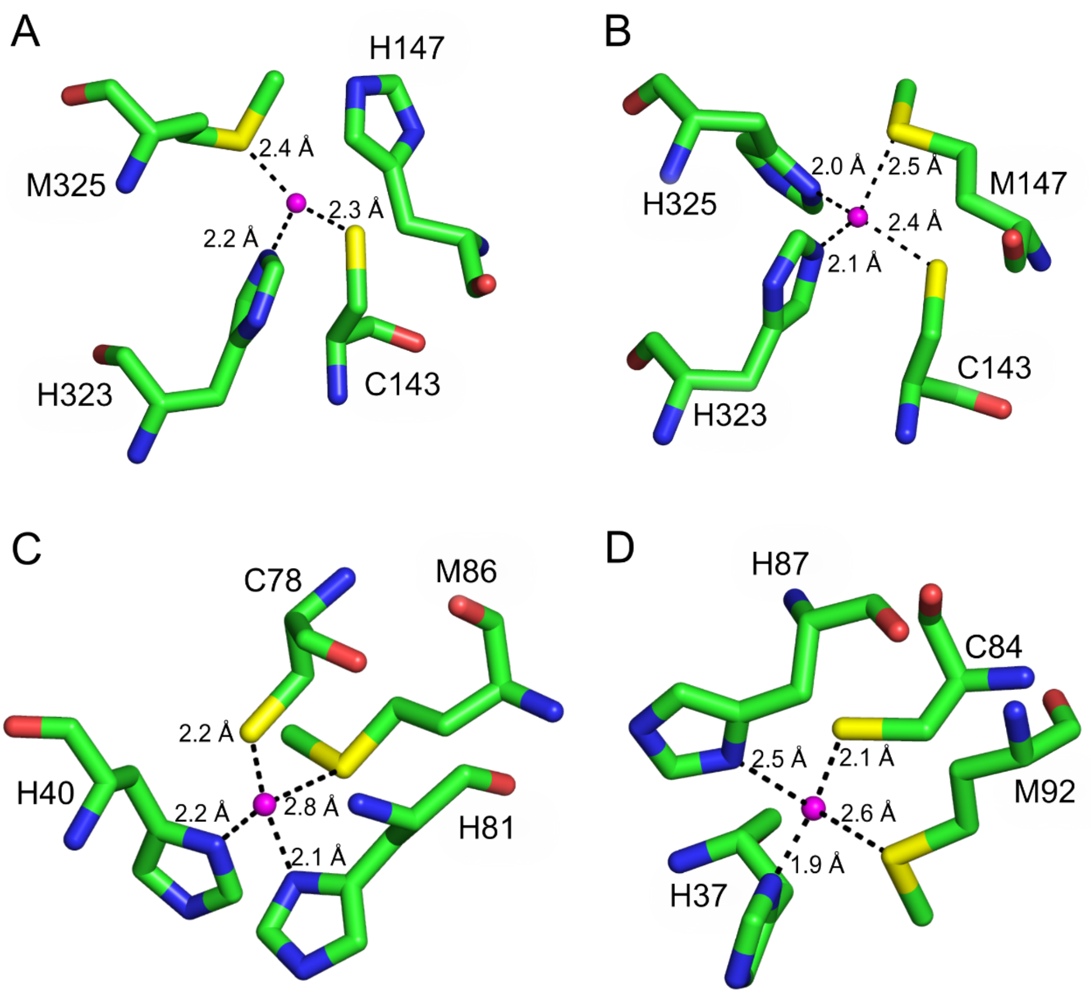


| **Bond distances (Å)** | **Pseudoazurin (1PAZ)** | **Plastocyanin (4DPA) Cu (I)** | **Plastocyanin (4DP9) Cu (II)** | **M147H** | **M325H** |
|---|---|---|---|---|---|
| Cys | 2.16 (Cys78) | 2.15 (Cys84) | 2.16 (Cys84) | 2.32 (Cys143) | 2.43 (Cys143) |
| Met | 2.76 (Met86) | 2.62 (Met92) | 2.78 (Met92) | 2.40 (Met325) | 2.5 (Met147) |
| His | 2.16 (His40) | 1.91 (His37) | 1.94 (His37) | 2.16 (His323) | 2.08 (His323) |
| His | 2.12 (His81) | 2.54 (His87) | 1.99 (His87) | - (M147H) | 1.99 (Met325) |
| **Geometry (*)** | Tetrahedral | Tetrahedral | Tetrahedral | Trigonal planar | Tetrahedral |

(*) Confirmed by the CheckMyMetal server (Zheng et al, 2014)

Figure S7. Comparison of M147H and M325H binding site residues with pseudoazurin and plastocyanin. Close up views of copper binding site residues in (A) M147H, (B) M325H, (C) pseudoazurin (PDB ID 1PAZ) and (D) plastocyanin (PDB ID 4DPA). The bound form of copper is Cu (I). Distances between coordinating residues and metal (magenta) are shown. The Table summarises distances between copper and co-ordinating residues as well as geometry.

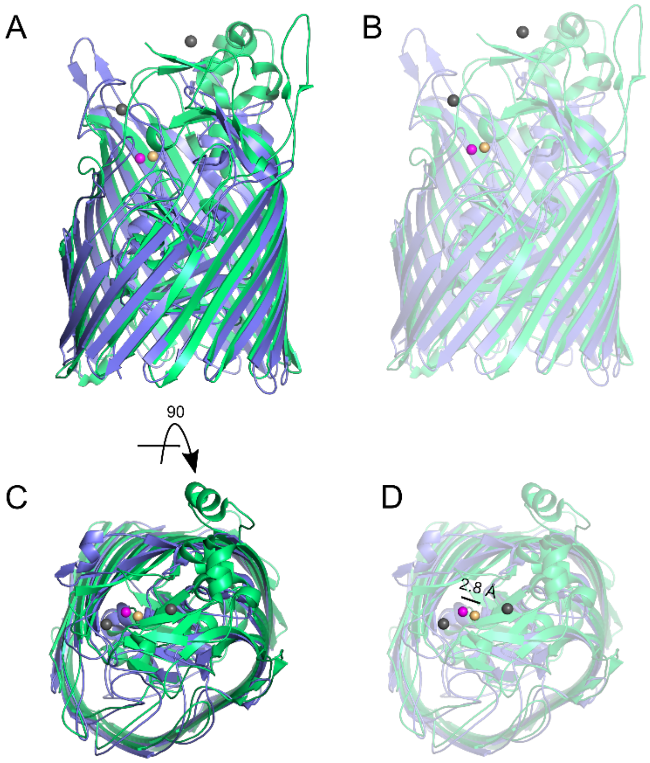


Figure S8. Differences between OprC and the Zn-specific ZnuD. (A, C) Cartoon representation comparing Cu-loaded OprC (coloured blue, copper atom shown as magenta sphere) and the locked version of ZnuD (coloured green, 2 cadmium atoms bound to low affinity sites represented as grey spheres, zinc bound to the high affinity site represented in orange; PDB ID 4RDR). The locked conformation of ZnuD shows low-affinity metal sites at external loops, at regions similar to the methionine track (L5) from OprC. (B, D) Transparent view of the secondary structures illustrates the similar topological location for the high affinity metal sites (distance of 2.8 Å).

Table S1. Data collection and refinement statistics for OprC variants with and without copper.

| | Cu - OprC | $OprC_{AA}$ | Cu – C143A 8800 | Cu – C143A 9175 | H323A – Cu 8800 | H323A – Cu 9175 |
|---|---|---|---|---|---|---|
| **Data collection**[#] | | | | | | |
| Space group | C 2 2 21 | P2 21 21 | P 2 21 21 | P 2 21 21 | P 1 21 1 | P 1 21 1 |
| Cell dimensions | | | | | | |
| *a*, *b*, *c* (Å) | 156, 197, 166 | 62, 171, 198 | 171, 198, 67 | 171, 198, 67 | 67,198, 172 | 67, 197, 172 |
| α, β, γ (°) | 90, 90, 90 | 90, 90, 90 | 90, 90, 90 | 90, 90, 90 | 90, 89, 90 | 90, 91, 90 |
| Resolution (Å) | 84.67 – 1.97 | 98.02 – 2.77 | 198.04 -2.78 (2.83 - 2.78) * | 129.57 -2.56 (2.60 - 2.56) * | 197.67 -2.95 (3.00 -2.95) * | 197.27 -2.73 (2.78 -2.73) * |
| $R_{pim}$ | 0.062(0.842) | 0.041 (0.748) | 0.078 (1.019) | 0.096 (2.48) | 0.081 (0.554) | 0.056 (0.577) |
| $I / \sigma I$ | 15.8 (1.3) | 4.4 (1.6) | 5.52 (0.39) | 5.40 (0.35) | 5.56 (0.58) | 7.82 (0.57) |
| $CC_{1/2}$ | 0.981 (0.784) | 0.990 (0.56) | 0.994 (0.400) | 0.974 (0.338) | 0.992(0.551) | 0.997 (0.674) |
| Completeness (%) | 99.5(100) | 88.8 (71.1) | 100(100) | 100(100) | 100(100) | 100(100) |
| Redundancy | 25.4 (25.4) | 5.7 (5.5) | 12.7 (12.1) | 12.3(12.3) | 6.6(6.2) | 6.6 (6.4) |
| **Refinement** | | | | | | |
| Resolution (Å) | 84.67 - 1.97 | 48.67 - 2.9 | 64.66 - 2.78 | 67.02 - 2.56 | 64.81-2.95 | 64.84 - 2.73 |
| No. reflections | 178812 | 42223 | 58104 | 74371 | 93985 | 118703 |
| $R_{work}$ / $R_{free}$ (%) | 21/ 23.2 | 21.6 / 27.1 | 20.9/26.7 | 24/29.8 | 24.2 /30.3 | 21.7 / 26.7 |
| No. atoms | | | | | | |
| Protein | 10050 | 10058 | 10119 | 10148 | 20154 | 20221 |
| Water | 556 | - | 23 | 39 | - | 67 |
| *B*-factors | | | | | | |
| Protein | 49.27 | 50.16 | 70 | 72 | 67 | 64 |
| Water | 49.28 | - | 54 | 60 | - | 54 |
| R.m.s. deviations | | | | | | |
| Bond lengths (Å) | 0.008 | 0.010 | 0.009 | 0.008 | 0.009 | 0.010 |
| Bond angles (°) | 1.26 | 1.54 | 1.12 | 1.08 | 1.17 | 1.15 |

[#] One crystal was used for each data collection.
* Values in parentheses are for highest-resolution shell.

Table S2. Data collection and refinement statistics for OprC variants with silver.

| | Ag – OprC 8000 eV | Ag – C143A 8800 eV | Ag – C143A 9175 eV | Ag – H323A 8800 eV | Ag – H323A 9175 eV |
|---|---|---|---|---|---|
| **Data collection**# | | | | | |
| Space group | C 2 2 21 | C 2 2 21 | C 2 2 21 | C 2 2 21 | C 2 2 21 |
| Cell dimensions | | | | | |
| *a*, *b*, *c* (Å) | 156, 195, 167 | 154, 195, 165 | 155, 196, 166 | 156, 196, 166 | 155,196, 165 |
| α, β, γ (°) | 90, 90, 90 | 90, 90, 90 | 90, 90, 90 | 90, 90, 90 | 90, 90, 90 |
| Resolution (Å) | 121.98 - 2.71 (2.76 - 2.71) * | 165.47 - 2.60 (2.64 - 2.60) * | 84.18 - 2.68 (2.73 - 2.68) * | 70.41 - 2.86 (2.91 - 2.86) * | 63.14 - 2.61 (2.66 - 2.61) * |
| $R_{pim}$ | 0.028 (0.503) | 0.055 ( 0.630) | 0.056 (0.654) | 0.207 ( 3.005) | 0.182 (2.940) |
| $I / \sigma I$ | 20.12 (1.47) | 7.56 (1.11) | 7.64 (1.09) | 4.18 (0.34) | 5.00 (0.43) |
| $CC_{1/2}$ | 0.999 (0.653) | 0.992 (0.644) | 0.996 (0.575) | 0.992 (0.284) | 0.993 (0.317) |
| Completeness (%) | 99.9(100) | 99.9(97.6) | 99.9(96.7) | 98.2(97) | 98(96.9) |
| Redundancy | 35.5 (31.6) | 13.5 (13.8) | 13.5(13.9) | 13.1 (13.5) | 13.3(13.6) |
| **Refinement** | | | | | |
| Resolution (Å) | 97.64 - 2.71 | 82.74 - 2.6 | 82.81 - 2.68 | 70.41 - 2.86 | 60.78 - 2.61 |
| No. reflections | 69472 | 76648 | 70402 | 54313 | 73192 |
| $R_{work}$ / $R_{free}$ (%) | 21.6 / 25.4 | 22.2 / 26.1 | 21.4 / 25.2 | 20.7 / 27.2 | 21.4 / 27.4 |
| No. atoms | | | | | |
| Protein | 10054 | 10156 | 10182 | 10073 | 10112 |
| Water | 13 | 13 | 8 | - | 71 |
| *B*-factors | | | | | |
| Protein | 79 | 72 | 72 | 69 | 61 |
| Water | 63 | 57 | 60 | - | 52 |
| R.m.s. deviations | | | | | |
| Bond lengths (Å) | 0.008 | 0.008 | 0.009 | 0.009 | 0.008 |
| Bond angles (°) | 1.06 | 1.08 | 1.12 | 1.2 | 1.04 |

### One crystal was used for each data collection.

* Values in parentheses are for highest-resolution shell.

Table S3. Data collection and refinement statistics for M147H and M325H variants.

| | Cu – M147H 9175 eV | Cu – M325H 9175 eV |
|---|---|---|
| **Data collection**# | | |
| Space group | C 2 2 21 | C 2 2 21 |
| Cell dimensions | | |
| *a*, *b*, *c* (Å) | 155, 197, 165 | 156, 196, 167 |
| α, β, γ (°) | 90, 90, 90 | 90, 90, 90 |
| Resolution (Å) | 98.09 - 2.38 (2.42 - 2.38) * | 97.85 - 2.37 (2.41 - 2.37) * |
| $R_{pim}$ | 0.044 (1.056) | 0.044 (1.255) |
| $I / \sigma I$ | 11.71 (0.89) | 12.46 (1.00) |
| $CC_{1/2}$ | 0.996 (0.321) | 0.997(0.315) |
| Completeness (%) | 100 (100) | 100(100) |
| Redundancy | 13.6 (13.2) | 13.3(13.5) |
| **Refinement** | | |
| Resolution (Å) | 63.34 - 2.38 | 61.02 - 2.37 |
| No. reflections | 101122 | 103039 |
| $R_{work}$ / $R_{free}$ (%) | 21.8/ 25.3 | 21.9/ 25.2 |
| No. atoms | | |
| Protein | 10158 | 10164 |
| Water | 133 | 107 |
| *B*-factors | | |
| Protein | 65 | 66 |
| Water | 57 | 28 |
| R.m.s. deviations | | |
| Bond lengths (Å) | 0.009 | 0.008 |
| Bond angles (°) | 1 | 0.99 |

### One crystal was used for each data collection.
* Values in parentheses are for highest-resolution shell.
